# Cultivation, chemistry, and genome of *Psilocybe zapotecorum*

**DOI:** 10.1101/2023.11.01.564784

**Authors:** Dusty Rose Miller, Jordan Taylor Jacobs, Alan Rockefeller, Harte Singer, Ian M. Bollinger, James Conway, Jason C. Slot, David E. Cliffel

## Abstract

*Psilocybe zapotecorum* is a strongly blue-bruising psilocybin mushroom used by indigenous groups in southeastern Mexico and beyond. While this species has a rich history of ceremonial use, research into its chemistry and genetics have been limited. Herein, we detail mushroom morphology and report on cultivation parameters, chemical profile, and the full genome sequence of *P. zapotecorum*. First, growth and cloning methods are detailed that are simple, and reproducible. In combination with high resolution microscopic analysis, the strain was barcoded, confirming species-level identification. Full genome sequencing reveals the architecture of the psilocybin gene cluster in *P. zapotecorum,* and can serve as a reference genome for Psilocybe Clade I. Characterization of the tryptamine profile revealed a psilocybin concentration of 17.9±1.7 mg/g, with a range of 10.6-25.7 mg/g (n=7), and similar tryptamines (psilocin, baeocystin, norbaeocystin, norpsilocin, aeruginascin, 4-HO-tryptamine, and tryptamine) in lesser concentrations for a combined tryptamine concentration of 22.5±3.2 mg/g. These results show *P. zapotecorum* to be a potent – and variable – *Psilocybe* mushroom. Chemical profiling, genetic analysis, and cultivation assist in demystifying these mushrooms. As clinical studies with psilocybin gain traction, understanding the diversity of psilocybin mushrooms will assure that psilocybin therapy does not become synonymous with psilocybin mushrooms.

## Introduction

*P. zapotecorum* R. Heim emend G. Guzmán is a psilocybin producing basidiomycete fungus endemic to the neotropics. It was first described to science with a collection from the Zapotec region of present day Oaxaca, Mexico, where it is referred to as ‘derrumbe’, translating literally to ‘landslide’.^1–6^ Indigenous cultures of mesoamerica, particularly the Zapotec, Mixtec and Mazatec collect these mushrooms in the wet season from landslides, among other habitats, and consumed them in ceremony.^7–10^ Indigenous knowledge of its psychopharmacology is documented by pre-columbian ethnomycological glyphs, codices, and physical artifacts made from ceramic, stone, and metal.^1,2,10–14^ Yet it was not until 1956 that *P. zapotecorum* was described in the scientific literature. That year, the French mycologist Roger Heim created the first type description from a 1954-55 collection by Valentina Pavlovna Wasson and Robert Gordon Wasson, from Santiago Yaitepec, Oaxaca, Mexico. In 1957 it was described in greater detail from a collection by R. G. Wasson and R. Heim in Huautla de Jimenez, Oaxaca, Mexico, where they met Maria Sabina, a seminal figure in the merger of the scientific and traditional lineages of these mushrooms.^3–7^ In this work, we continue to build on the scientific knowledge of this fungus, while recognizing that deep relationships to it have already been forged.

Originally described as fruiting from swampy, muddy soils, *P. zapotecorum* is now known to fruit from multiple habitats and substrates. In Mexico, *P. zapotecorum* can be found in cloud forests at elevations between 900 and 3200 meters (**Figure 1**).^15^ It fruits abundantly in landslides, areas where water has carved out clay ravines, and anthropogenic disturbances such as land cuts for road development.

**Figure 1.**
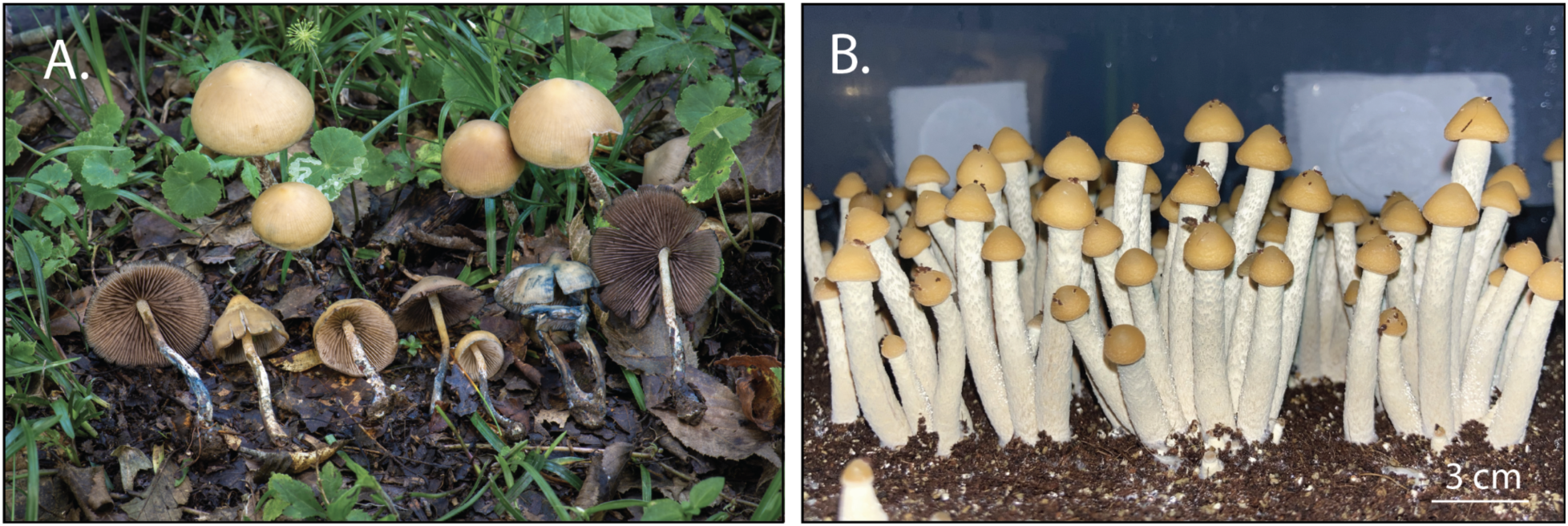
Photograph of wild (**A**) and cultivated (**B**) *P. zapotecorum*. Photograph of the wild collection was provided by Alan Rockefeller showing *P. zapotecorum* growing in a grassy area around La Martinica, Mpo Banderilla, Veracruz, Mexico at 19.5835°N 96.9485°W 1581m. Photograph of cultivated *P. zapotecorum* (**B**) showing fruits growing under defined conditions.

*P. zapotecorum* fruit bodies are somewhat polymorphous both macroscopically and microscopically, complicating species boundaries. In the past, *P. zapotecorum* was taxonomically split into several species, which have now been recombined. Since 1956, ten synonymous species and three varieties have been described based on slight morphological or habitat distinctions from the original *P. zapotecorum* description.^1^ An emendation of *P. zapotecorum* by examination of microscopic features and holotype reviews presents a new species concept, encompassing all of these species and varieties into *P. zapotecorum*.^1^

Ultimately, boundaries between species can vary depending on how the lines are drawn and molecular tools can aid in the process of both establishing and sorting within species boundaries. Full genome sequencing provides the highest phylogenetic resolution, while isolated DNA barcoding can often provide clear and reproducible evidence of speciation when either the genome sample, financial resources, or time is limited. Five genetic barcode loci are commonly used for fungi: internal transcribed spacer 1 and 2 and the intervening 5.8s ribosomal RNA gene (ITS), elongation factor 1-alpha (ef1-alpha), nuclear large subunit ribosomal RNA gene (nLSU), and RNA polymerase II largest and second largest subunit genes (*rpb1* and *rpb2*, respectively). Of the three barcode sequences previously determined for *P. zapotecorum* and well represented among *Psilocybe* in GenBank, ITS and *rpb1* were shown to be most informative for species delimitation.^16^

Contiguous genome assembly can assist in answering a broad range of questions, including resolving genetic diversity in secondary metabolite production and inferring genetic relatedness among species.^17^ Analyzing the complete genetic code enables prediction of the mechanisms that govern its growth, development, and psychedelic properties of fungal species.^18–20^ Furthermore, genome sequencing enables comparison of the genetic makeup of closely related species of fungi.^18,19^ Such comparisons paired with evolutionary analysis can provide insight about the unique adaptations of species like *P. zapotecorum*. A full genome of *P. zapotecorum* will be a valuable resource for future research into its therapeutic and medicinal applications.

Based on genetics, morphology and habitat, evolutionary relationships between *Psilocybe* species have been established. In 1983, a psilocybe monograph was published inferring genetic relationships based primarily on morphology and grouped psilocybe mushrooms into 18 sections with *P. zapotecorum* being contained within the eponymous section Zapotecorum.^21^ Species within section Zapotecorum are characterized by thin-walled subellipsoid, subrhomboid or subfusoid spores, hyaline or brown pleurocystidia (sometimes absent), and occur in subtropical or temperate climates. Subsequently, two monophyletic clades, clade I and clade II were proposed based on sequencing multiple loci of *Psilocybe sensu stricto*. ^22^ Within this organization, the clades are split into monophyletic subclades A & B within clade I, and C & D within clade II. *Psilocybe zapotecorum* is placed within clade I, subclade A. Subclade A contains neotropical species sharing habitat and morphological characteristics, while species in subclade B grow in temperate climates. Clade I contains species predominantly found in the Americas, whereas Clade II contains both temperate and tropical species of *Psilocybe* with cosmopolitan distribution and includes the commonly cultivated species *Psilocybe cubensis*. While *P. cubensis* does occur in the neotropics, it may have been unintentionally introduced during the Spanish conquest with the introduction of cattle ranching. Unlike *P. cubensis*, *P. zapotecorum* is endemic to the neotropics and resides within Clade I, subclade A alongside section Mexicanae.

Despite its wide-spread distribution *P. zapotecorum* has proven difficult to cultivate and its use has been limited to wild-gathered mushrooms. Although *P. zapotecorum* were some of the first *Psilocybe* mushrooms cultured, they are rarely cultivated. The reason for this may be due to the long growth period reported by Heim, or the difficulty of rearing fruit bodies to maturity. The first reported cultivation of *P. zapotecorum* in the scientific literature was by Roger Heim in 1957, fruiting the fungus in a glass flask with a few different mixes including sterilized moss, compost, or straw, sometimes mixed (or cased) with sand.^4^ To our knowledge, only one other instance of successful cultivation of *P. zapotecorum* has been published in the scientific literature.^23^ In contrast, there is a wide body of knowledge available regarding the cultivation of *P. cubensis*– it grows rapidly and is amenable to variable environmental conditions. This is reflected in the literature: with most of the information on psychedelic mushrooms limited to this species it creates a foundational – albeit skewed – perspective of psilocybin fungi.

A key feature of psilocybin containing mushrooms is a characteristic blue bruising of the flesh upon handling. In *P. zapotecorum*, the blue pigment materializes as royal electric blue, rapidly intensifying to a charcoal black, earning the mushroom the common name *Derrumbes Negro.* This trauma-induced bluing is a enzymatic reaction that’s’ mechanism has been characterized in detail in *P. cubensis*.^24,25^ With regards to *P. zapotecorum*, it is not known whether the color shade and intensity observed is due to higher concentrations of psilocybin, higher activity of psilocin oligomerization enzymes, or multiple compounding factors.

Tryptamine concentrations do not appear to be elevated compared to other psilocybin containing mushrooms [Stijve and de Maijer (0.6-3 mg/g psilocybin, 0.5-10 mg/g psilocin, up to 0.2 mg/g baeocystin), Heim and Hofmann (0.5 mg/g psilocybin, 0 mg/g psilocin, 2 year old specimens)].^26,27^ However, it has been shown that psilocybin concentrations are highly variable, can decrease over time, and are dependent on storage conditions.^28–30^

When ingested, the effects of psilocybin mushrooms can vary but have been shown to treat psychological disorders and occasion mystical experiences. Psychedelics are known for their changes in perception including time dilation or contraction, auditory intensification or dampening, rhythmic enhancement, and open- or closed-eye visions. Although it is not well understood how each of these perceptual changes arise, brain imaging scans indicate the importance of the default mode network in decreasing typical or repetitive circuitry, allowing new connections to form.^31^ There may also be changes in feeling states or perspectives marked by immense laughter, comfort, profound insight, and sensations of love and connection.^32–34^ These experiences can have lasting impacts on life perspectives, beliefs and feelings of well-being and have been shown to treat depression, anxiety including end of life anxiety, post-traumatic stress disorder, and alcohol use disorder among others.^32–38^ In treating these conditions with psychedelics, it is clear that set and setting have a profound impact on the experience.^39,40^ Research into another psychedelic, ketamine, highlights a biochemical explanation for its healing attributes – resetting of the homeostatic or baseline neuronal signaling.^41,42^ New research into lysergic acid and psilocybin suggests that there may be avenues for healing outside of serotonergic pathways, such as activation of brain derived growth factor.^43^ With *P. zapotecorum*’s widespread use by indigenous peoples, it begs the question, is it merely what is available, or does this mushroom have distinct qualities?

There are many psychoactive fungi but legislation is not treating them the same. There are thought to be around 200 different species of psilocybin mushrooms, each with distinct qualities and chemistries.^44^ Oregon voters recently legalized psilocybin mushrooms, initiating a cutting-edge program where people can ingest psilocybin mushrooms under the guidance of a facilitator.^19,45^ However, only *P. cubensis* is permitted for use. As clinical studies of psychedelic-assisted psychotherapies gain traction, it is important to continue investigating the naturally occurring source of psilocybin in mushrooms, so that synthetic psilocybin, used in most clinical studies, or *P. cubensis*, used in Oregon, does not become unintentionally synonymous with mushroom therapy. This work may provide scientific evidence to support the safe inclusion of *P. zapotecorum* in therapeutic use.

The brilliant bluing combined with the historic use of this mushroom suggest it is worthy of further investigation. This publication expands the current scientific understanding of *P. zapotecorum,* by describing strain development, cultivation, microscopy, and genetic and chemical analysis. Starting from wild-collected spores, a strong fruiting strain was found, isolated, and cloned. Using common substrates or industrial byproducts, a simple and robust cultivation method was developed, allowing for careful collection and rapid processing. We used microscopy to show the micromorphological features in detail. High performance liquid chromatography (HPLC) of cultivated basidiomata revealed a new perspective on *P. zapotecorum* tryptamine alkaloid content. Full genome sequencing also provides insight into the psilocybin gene cluster and evolutionary relationships. This multidisciplinary approach adds some distinction to the chemistry of this revered mushroom species and provides a strain for continued investigation. This study is poised to support clinical investigations into the therapeutic potential of psilocybin mushrooms and provide evidence for the safe inclusion of *P. zapotecorum* in legislation where these novel therapies are being explored. Continued investigation of these mushrooms will help normalize them both as medicines and as entheogenic sacraments used by indigenous peoples and sincere churches.

## Materials and Methods

### 1.1 Fungal cultivation

*P. zapotecorum* was grown through a complete life cycle with an intermediate process targeting the establishment of a basidome-producing clone. Briefly, wild spores were streaked as multispore plates and left to grow basidiomes. A somatic fragment of a basidiome was removed, plated and allowed to expand. Actively growing mycelial material was removed and transferred to liquid culture (LC). This LC was used for fruit production by combining it with either grain or wood, allowing it to colonize, and then mixing this spawn with substrate. During fruit growth, humidity, aeration, and lighting were provided. These growth conditions below detail the steps for *P. zapotecorum* to be taken through a complete life cycle – from spore to spore.

Spores were collected by Alan Rockefeller and Alonzo Cortes-Perez from basidiomata found fruiting near Xalapa, Vera Cruz, Mexico. Spores were purchased from Alonzo Cortes-Perez and germinated in Petri dishes on malt extract agar [MEA, 15 g/L malt extract, (Breiss, Chilton WI), 1 g/L yeast extract (Shroom Supply, Brooksville FL), 18 g/L agar (Shroom Supply), Elkay filtered municipal water (pH unadjusted)] and maintained on the same medium at 20-22°C.

The multispore plate was subcultured for diverse basidiome formation. As mycelial growths emerged from the multi-spore plate, they were removed in large 20mm^2^ sections and replated to create a diverse set of mycelium. These subcultures were allowed to grow at room temperature for 90 days.

*In vitro* basidioma formation was observed, the basidioma was cloned, and its mycelium was expanded. Cloning was achieved by aseptically extracting the basidioma from the Petri dish, tearing the stipe open to expose sterile clonal inner tissue, taking a small explant of the inner tissue, and plating the explant on a new MEA Petri dish. The strain resulting was named ‘*La Martinica*’ to represent the locality of the collection. The mycelium from this explant was allowed to colonize the Petri dish to 70%. Then the clonal mycelium was expanded by taking a 5mm^2^ square of mycelium from the leading edge of the colony and replating it. This culture was placed in the fridge for storage except for a small fragment that was used to create a liquid culture.

The La Martinica LC was expanded for various purposes and/or maintained. This was done by inoculating a malt extract liquid medium (LM,3% malt extract, 0.3% yeast extract (w/v%, Shroom Supply)] with a 5mm^2^ portion of mycelium from the Petri dish. The LM was stirred continuously at 140 rpm for 14 days at 20-22 °C, yielding LC. To expand the mycelium, fresh LM was inoculated with LC at a 1% (v/v) inoculum to medium ratio. After another 14 day expansion, this LC was used to inoculate grain. To create mycelium for DNA extraction, mycelium that had been colonized for 3 weeks at room temperature was added to an autoclaved stainless steel blender jar and homogenized with sterile water before being added to 2% malt extract yeast peptone broth where it was shaken at room temperature for 17 days. This mycelium was then harvested and the DNA was extracted (see section **1.5 Genome sequencing and analysis**).

Grain spawn was prepared by inoculating sterile grains with LC. Whole oats were hydrated in boiling water until the point of cracking, then removed from the heat and spread out to dry. When cool, they were added in 1kg portions to mushroom spawn bags (Shroom Supply). The air was pressed out of the remaining volume in the bag and the bag was folded. Bags were autoclaved at 121°C for 1.5 hours to sterilize. The sterilized grain bags were inoculated with LC at 0.5% (v/wt) inoculum, then sealed using an impulse sealer. The grain bags were shaken to distribute inoculum, then incubated in the dark at room temperature. When the myceliated grain in the bag was between 30% to 70% colonized, the myceliated areas were broken up and the grain was mixed. When the grain was 100% colonized, it was considered grain spawn and suitable for transfer to a fruiting chamber.

To provide proper fruiting conditions, a simple fruiting chamber was constructed. The fruiting chamber was based on designs utilized for fruiting *P. cubensis*. The chamber allowed for light penetration, fresh air, and humidification. The chamber was made from a clear plastic storage bin (Sterilite, Townsend, MA) that was modified by boring six 50mm holes and covering these holes with layers of breathable medical micropore tape (3M, Maplewood, MN). Once the bin was modified, a plastic liner was inserted, and the chamber was ready to be filled with substrate.

To encourage fruiting, spawn was mixed with substrate and added to the fruiting chamber. A substrate designed to hold water and allow air flow was made. Coconut coir vermiculite substrate, hereafter called ‘substrate’ was prepared from coconut coir, vermiculite, and water by combining 3.8L of 80 °C water with the dry ingredients (650g coconut coir, 2 quarts vermiculite), then mixing until homogeneous. The substrate was prepared to substrate at field capacity (maximum water holding capacity) and allowed to cool to room temperature before use. Spawn was mixed with substrate in the fruiting chamber at a ratio of 1:2 (w/w), respectively. Once evenly mixed, a 1.5 cm top layer of substrate was applied and the bin was ready to connect to the humidifying chamber.

To maintain ideal humidity in cultivation, a humidification chamber was constructed. The humidifier was made from an ultrasonic fogger, fogger float, and a waterproof fan (all Shroom Supply) inside a dedicated clear storage bin. Two holes were drilled in the lid of the bin that were the diameter of the fan. The fan was fixed to one hole, and a flexible duct was fixed to the other. An additional hole was drilled into the lid for the power supply cord of the fogger. To operate, the tote was filled with water, and the fogger floated in the water. The fan and fogger were turned on and fog flowed from the tote through the flexible ducting to the fruiting chamber, supplying mist. The fruiting chamber was fogged for about 15 minutes once a day and visually inspected. Indication of proper humidity was the presence of residual water droplets on the interior surfaces of the fruiting chamber. Water was not allowed to pool on the surface of the substrate. Humidity was maintained through fruit production.

Once mature, basidiomata were harvested, dried and stored for further use. Basidiomes were harvested using scissors to sever them at the base and were carefully placed on drying racks and immediately set in a dehydrator (Ivation, Edison, NJ). The basidiomata were then dried at 37-39°C and either analyzed immediately or stored in an airtight container with moisture and oxygen absorbers for later analysis.

### 1.2. Microscopy

Microscopy images of spores, fruit body cross sections and gill fragments were obtained and edited with post-acquisition processing.

Cultivated and freshly dried La Martinica fruit bodies were dissected for microscopy. Razor dissection was performed by hand under an Amscope stereo zoom microscope (United Scope, Irving California, USA) and dissection razors were used for less than 10 slices before being discarded. To measure the spores, a thin scalp section of pileipellis with spore deposit was taken. The lamella edge was prepared from an excised lamella fragment. Cross sections of the pileus were prepared for pileus and lamellae inner tissue. The scalp section, gill fragment, and thinnest cross section were cut so that the final size of the sections was less than 0.25mm^2^. The dissected scalp section, lamellae fragment, and cross sections were placed on individual microscope slides (United Scope) and rehydrated in 5% potassium hydroxide (wt/v) for two minutes. High precision coverslips #1.5H (Thor Labs) were applied, and gently pressed down to displace air bubbles. Excessive moisture, if present, was wicked away using tissue paper. Coverslips were adhered in place using nail polish to prevent evaporation before being imaged.

Microscope slides were viewed using an Olympus BX41 compound microscope (Olympus-lifescience, Tokyo, Japan) in differential interference contrast viewing mode. Magnification was achieved with 40x and 100x Uplanapo oil immersion objectives. Images were taken using a Nikon Z7 camera body (Nikon, Tokyo, Japan) mounted to the trinocular head. A remote shutter release was utilized to minimize vibrations during image capture. Images were taken in series with successive focus adjustments (focus shift) to capture the entire depth of field.

Images of the microscopic subjects at varying focal points were stacked to achieve full focus images. Image stacking was performed in Helicon focus 8 (Helicon Soft, Kharkiv, Ukraine). Images were further processed in Photoshop (Adobe, San Jose, CA). Adjustments of color levels were made to achieve images with more accurate color profiles. Spore measurements were taken using Piximètre (version 5.1, revision 1541).

### 1.3. DNA barcoding

Species authentication was performed by sequencing the Internal Transcribed Spacer (ITS) region of rDNA from the La Martinica strain. To isolate DNA for PCR amplification, approximately 500μg of tissue was taken from a culture of *P. zapotecorum* grown on malt extract yeast agar [Malt Extract Broth (Sigma-Aldrich) 20g/L, Yeast Extract (Shroom Supply) 2g/L, Agar-Agar powder 20g/L (Landor Trading Company, China)] and added to a 200μL PCR tube using flame-sterilized tweezers in a laminar flow cabinet. Then, 20μL of X-Amp DNA Reagent (IBI Scientific, Peosta, IA) was pipetted over the sample and the sample was disrupted for 30 seconds using a sterile disposable plastic pestle. The PCR tube was incubated at 80 °C for 10 minutes before cooling to 12 °C using a GeneAmp 2400 thermal cycler (Perkin Elmer, Waltham MA).

The nrITS sequence from the *P. zapotecorum* DNA isolate was amplified using PCR. DNA isolate (2μL) was added directly to a 25μL PCR reaction with 8.5μL molecular grade water, 12.5μL 2x Taq PCR master mix (Dikarya, Santa Rosa CA) and 1μL each of ITS1F and ITS4 primers (10μM, Dikarya) and run on a GeneAmp 2400 (Perkin Elmer) thermal cycler with the following conditions: First a 95 °C hold for 4 minutes, followed by 35 cycles of 95 °C for 30 seconds, 55 °C for 30 seconds, 72 °C for 1 minute, then followed by a final extension and hold of 72 °C for 7 min and rested at 12 °C.

DNA amplification was confirmed with gel electrophoresis. A 1% agarose gel stained with GelRed® Agarose LE (Biotium, Fremont CA) in Borax buffer [1g/L Sodium Tetraborate (20 Mule Team, Rocky Hill CT)] was run for 15 minutes at 200V. Visual confirmation of a band indicated successful amplification.

After successful amplification, 10μL of PCR product was sent to Genewiz (Azenta Life Sciences, South San Francisco, CA) for Sanger sequencing using ITS1 and ITS4 primers. The electropherograms were analyzed and a contiguous sequence was constructed using Geneious Prime 2023.0 (Biomatters, Boston, MA). The consensus sequence was used in phylogenetic analysis.

### 1.4 Analytical chromatography

A sample preparation protocol by Dorner et al. 2022 was followed with some modification to analyze tryptamine alkaloids.^46^ Dried carpophores were kept whole or separated by cap and stem for a comparative analysis. The portions were commutated with a mortar and pestle to a fine powder. Then, 30.0mg ± 1.0mg portions of homogenate were added to 2mL centrifuge tubes. 1mL of methanol was added to the tubes, then the tubes were sealed. The samples were vortexed, then placed in an ultrasonic bath and continuously sonicated for 10 minutes at room temperature. The samples were then centrifuged for 5 minutes at 14,000 rpm. The supernatant was collected, and 1mL of methanol was added to the tissue pellet for a second extraction following the same conditions as the first. The second supernatant was combined with the first, and a third extraction was performed on the tissue pellet with 1mL of a 70% methanol and 30% water with 0.1% formic acid (FA) mixture. Besides this change in solvent the third extraction was performed the same as the first two. The third supernatant was pooled with the first two and filtered into an autosampler vial using a 0.22um PTFE filter(Biomed Scientific, Los Angeles, CA). Samples were run as is (for minor tryptamines) and diluted 1:10 (for psilocybin). Samples were either immediately analyzed or stored at −20 °C for later analysis.

Analytical HPLC was performed on an Accela 600 HPLC equipped with an Accela PDA detector (Thermo Fisher). Separation of tryptamine alkaloids in mushroom extracts was achieved with a biphasic gradient elution on a Kinetex® Biphenyl 5 *μ*m 100 Å 150 mm x 3.0 mm LC column (Phenomenex, Torrance, CA) and a EXP guard column (Restek, Bellefont, PA). The sample injection volume was set to 0.5 μL. The column compartment was held at 26°C, and the flow rate was held at 0.75mL per minute. Mobile phase A was HPLC grade water fortified with 0.1% FA (V/V %), and mobile phase B was neat HPLC grade methanol. Initial conditions were 97.5:2.5 A:B, and the gradient elution method was as follows: hold initial conditions from 0.0-0.5 min, to 90:10 by 1.0 min, to 80:20 by 4.0 min, to 70:30 by 5.5 min, to 0:100 by 10.0 min, and back to initial conditions by 10.01min and held there until 15 min. The detection and quantitation wavelength was set to 280.0nm, and the peaks matching retention times of certified reference materials were scanned with the DAD from 190-450 nm for identity confirmation.

Analyte quantitation was performed using multipoint calibration with a linear equation from peak area. Starting from 1 mg/mL solutions of certified reference materials (Cayman Chemical Company, Grand Rapids, MI), 100uL of each analyte was added to the standards mix vial, then diluted with methanol to 1mL, achieving a concentration of 100μg/mL. Serial dilutions of this 100μg/mL mixture were performed to achieve standard concentrations of 50, 25, 10, 5, and 1 μg/mL. The standards at various concentrations were run on the HPLC with all instrument parameters consistent with sample runs. The peak area for each analyte at each calibration level was added to the calibration table. When analytical runs were performed, samples were bracketed with continuous calibration verification (CCV) controls for accuracy and precision of analysis.

The limits of detection (LOD) and limits of quantitation (LOQ) for the method were estimated using low level CRM spiked in background and matrix matched blanks. For the background blank for the study a non-psilocybin containing mushroom, *Agrocybe pediades* was selected, as it did not contain any compounds matching the retention times of the quantified tryptamines. This was homogenized according to the *P. Zapotecorum* protocol was and 30 mg of this homogenate was weighed in a 2mL extraction tube. Then, 6 uL of 100 ug/mL CRM mix was spiked onto the homogenate and allowed to saturate into the matrix blank for 30 minutes at room temperature under ambient light. The spiked sample was then extracted, prepared, and analyzed as above. Then, 10 replicate extractions were performed, and each sample was analyzed once with an expected final concentration of 0.2 ug/mL. The standard deviation of each analyte peak area for the 10 replicates was used to calculate the LOD and LOQ. The LOD and LOQ for the method were estimated as three and ten times the standard deviation of analyte peak area respectfully.

### 1.5 Genome sequencing and analysis

To determine the genetic code of *P. zapotecorum* and provide a reference for future work, the full genome was sequenced. First sufficient genetic material was harvested (see section **1.1 Fungal cultivation**), and DNA was isolated. To isolate DNA the harvested mycelium was strained, frozen in liquid nitrogen, and crushed with a mortar and pestle and the DNA was isolated using the Qiagen DNeasy ® Plant Mini kit following the manufacturer’s instructions for high molecular weight DNA. DNA was quantified using the Qubit™ 4 Fluorometer, and DNA integrity was confirmed using a 2% agarose gel stained with GelRed ® and DNA purity was checked with Nanodrop 6000 (Thermo Fisher). The gDNA generated was sent to Novogene, Inc (Novogene USA, Sacramento, CA) where both Oxford Nanopore Technologies (ONT) and Illumina library preparation and sequencing were conducted. Long reads were generated using the ONT PromethION sequencing platform on a Flow Cell R9.4.1 using Ligation Sequencing Kit SQK-LSK110, and base-called with Guppy v6.4.6 HAC High accuracy. The Illumina data was generated using a NovaSeq 6000 system with paired-end 150-bp reads.

The Oxford Nanopore Technologies (ONT) sequences were processed to result in an indexed partial assembly. Long reads were generated using the ONT PromethION sequencing platform on a Flow Cell R9.4.1 using Ligation Sequencing Kit SQK-LSK110, and base-called with Guppy v6.4.6 HAC High accuracy. Unprocessed reads were de novo assembled using Flye v.2.9.2 creating an assembly with 1180 unique contigs.^47^ The initial Fly assembly was checked for contamination through BLASTn searches, filtering out contigs by looking at three sections of each individual contig over 500bp and separating non-fungal matches with average Percent ID: >95%. The filtered Flye assembly was then indexed with the BWA-MEM algorithm. This indexed assembly was then ready to combine with the trimmed Illumina reads.^48^

Paired-End Illumina reads were trimmed using Trimmomatic v0.39 with the following parameters: PE -phred33, ILLUMINACLIP:TruSeq3-PE.fa:2:30:10:11, LEADING:25, TRAILING:25, SLIDINGWINDOW:50:20, and MINLEN:125.^49^ Quality of trimmed reads was assessed using FastQC (v0.11.9).^50^. Unprocessed ONT reads were assembled *de novo* using the Geneious Prime (Version 2023.1.2, Biomatters, 2023, available from https://www.geneious.com) plugin for Flye (v.2.9.2 de novo assembly creating an initial assembly with 1180 unique contigs.^47^ This assembly was checked for contamination through BLASTn searches, filtering out contigs by looking at three sections of each individual contig over 500bp and separating non-fungal matches with average Percent ID: >95%. The filtered Flye assembly was then indexed with the BWA-MEM algorithm for polishing with the trimmed Illumina reads.^48^ Trimmed paired-end Illumina reads were aligned to the ONT Flye Assembly using the Burrows–Wheeler (BWA-MEM) algorithm resulting in an alignment stored in a Sequence Alignment/Map (SAM), which was converted into a Binary Alignment/Map (BAM) format, and sorted and indexed based on the reference sequence coordinates using SAMtools.^51^ Finally, the assembly was refined using Pilon, treating the organism as diploid.^52^ The final Flye assembly was assessed using the Quality Assessment Tool for Genome Assemblies v5.2.0 (QUAST) and benchmarked by running the Benchmarking Universal Single-Copy Orthologs (BUSCO) v5.4.7 against the Basidiomycota and Agaricales databases (agaricales_odb10 and basidiomycota_odb10, available from https://busco-archive.ezlab.org/data/lineages).^53,54^

Psilocybin gene sequences (26) were obtained by first identifying the locus on the assembly that contains them with tblastn and then using exonerate to identify coding regions with reference to the *P. Mexicana* FSU_13617 assembly (GCA_023853805.1) and *P.* cyanescens (GCA_002938375.1) and P*. cubensis* (https://mycocosm.jgi.doe.gov/Psicub1_1/Psicub1_1.home.html) annotations.^55,56^ Psilocybin gene cluster loci for the four species were aligned in Clinker v.1.1.0.^57^

### 1.6 Phylogenetic analysis

The phylogenetic placement of *P. zapotecorum* La Martinica was analyzed by ITS and the RNA polymerase II largest subunit (*rpb1*) sequence in alignments containing sequences from clade I and clade II.^58,59^ Sequences of PCR-amplified ITS and *rpb1* derived from the full genome sequence were used to search NCBI using BLASTn and a selection of the BLAST results were downloaded for construction of the phylogenetic trees. *Rpb1* was annotated using a *Coprinopsis cinerea* reference sequence (KAG2023660.1) with the exonerate software package.^56^ Introns were removed from *rpb1* prior to analysis. Sequences of both genes were aligned using MAFFT (Version 7.4.07) and manually trimmed using the alignment viewer in the Geneious Prime software suite.^60^ The ITS tree was constructed under the TPM2u+F+G4 substitution model and the *rpb1* tree under the TNe+G4 substitution model, determined by the included model test in IQ-TREE (Ver 1.6.11).^61,62^ Support for topologies was assessed by 1000 ultrafast bootstrap replicates.

## Results

### 2.1 Fungal cultivation

*P. zapotecorum* was grown from spore to basidiome using various substrates at different points along the fungal life cycle. First, wild basidiomata spores (**Figure 1A**) germinated and growth was observed. These multispore culture plates grew and produced basidiomes. *In vitro* somatic clonal mycelium colonized liquid culture, grains, and substrate in a fruiting chamber. Fruitification was found to be reproducible.

Successful germination of wild *P. zapotecorum* spores was observed on MEA Petri dishes after ∼4 days. All of the resulting growths from subculturing of the initial spore germination plates displayed tomentose mycelial morphology. The mycelium was observed to have clamp connections under 400X magnification, indicating a dikaryotic mycelium had formed. A total of 20 spore inoculation plates were made, and the most vigorous three plates were selected and further subcultured to 20 new plates.

The subcultured dikaryotic plates produced basidiomata *in vitro*. After ∼9 weeks of incubation at 22 °C, basidiomata formed from one out of 20 subcultures in a Petri dish. The *in vitro* basidiomata developed to maturity and produced and ejected basidiospores. A single somatic clone of the largest basidioma was isolated and used in all subsequent experiments. This strain was named ‘La Martinica’. The clonal mycelium morphology was tomentose and did not change color or produce pigment in the media. Growth of the clonal mycelium was found to be sufficient with MEA at 20-22 °C. The culture continued to produce basidiomata *in vitro* when subcultures of its mycelium were left to incubate for nine weeks.

Mycelium grew in liquid and solid media. When liquid culture inoculum was introduced into fresh liquid media (1% v/v), the media was colonized after 10-14 days. Inoculation of liquid culture to whole oats (0.5% v/wt) showed visible growth within three days and full colonization within three weeks.

Using a fruiting chamber design based on those used for the fructification of *P. cubensis*, there was successful initiation of primordia and development into mature basidiomata (**Figure 1B**). Spawn to substrate ratios of 1:2-1:5 were sufficient to produce fruit bodies. During colonization, the fruiting chamber held proper humidity without any humidification. The mycelium colonized the substrate, except the casing layer. After complete colonization, hyphal knots formed, which developed into primordia. The first primordia formed 19 days after full colonization of the substrate, or 33 days after the fruiting chamber was spawned. Primordia originated from beneath the uncolonized substrate. Development of primordia into mature basidiomata continued over a period of 35 days from initial primordia observations. It was during this period of development that supplemental humidification promoted basidiomata formation. Successful fruiting required careful attention to humidity. When sprayed instead of fogged, fruits aborted. The caps of *P. zapotecorum* were hygrophanous-shiny when wet and opaque when dry. If the caps dried during cultivation, the fruits aborted. Fruits grew upward toward the light. Fruiting patterns were gregarious, forming dense clusters throughout the fruiting chamber. After 35 days, mature basidiospores could be seen ejecting spores from the hymenium of the basidiomata, leaving a purple deposit on the stipe, and eventually on the top of the pileus. The total time from inoculation of substrate to mature basidiomata was 68 days. When all these steps are done in direct succession, *P. zapotecorum* was taken from spore to spore in approximately 24 weeks. Starting with clonal LC, the process was approximately 13 weeks.

When the fruit bodies ejected spores, they were harvested and dried for analysis. Handling of basidiomata resulted in rapid pigment evolution at the site of trauma. It was found that basidiomata of the La Martinica strain had pseudorhiza that would pull up a clump of substrate when harvested. Because of this phenomenon, scissors were used to remove the mature basidiomata, leaving the pseudorhiza in the substrate.

*P. zapotecorum* La Martinica yielded uniform clusters of sturdy fruits with visible sporulation and variable sizes, resembling those found in the 2019 collection (**Figure 1 A-B**). Cultivated primorida initiated beneath uncolonized substrate, and developed into tight, caespitose clusters. Stems were decorated with floccose adornments, caps scalloped, finely papillate, with inrolled margins and partial veil remnants. Caps varied from umbonate, convex, or subsecotioid, and the ones that matured, ejected spores. Many of the developing mushrooms aborted in initial trials, before humidification intervals were defined. If left unharvested, the caps began to blacken. Sizes of spore-ejecting fruits varied, but grew as tall as 33cm with caps as wide as 8cm. On the inside, the stems were dense with woody brown mycelium and a hollow center channel.

### 2.2 Microscopic analysis

*P. zapotecorum* images were acquired of the pileus and spores (**Figure 2 A-D**). Microscopic images of *P. zapotecorum* show **A)** Spores, **B)** Gill edge, **C)** Pileus cross section, and **D)** Gill cross section. The spores were inequilateral symmetrical and ellipsoid to fusoid in shape, measuring averaging 6.3 ± 0.3 µm in length with a range of 5.9-6.9 µm and 3.8 ± 0.1 in width with a range of 3.6-4.1µm, n=29. Under microscopic investigation, color shows as golden with interspore variation in translucence, whereas by eye the spore color is purple to purple-brown. The spores showed truncated germ pores, an acute apiculus, and contained 3 ± 1 spherical lipid droplets of varying size, n=29. On the gill edge, both cheilocystidia and pseudocystidia can be seen. The cheilocystidia were hyaline, polymorphous, langeniform, ampulliform and sometimes furcate. Pseudocystidia were gray, polymorphous, sublageniform, subventricose, subpyriform, sometimes furcate and sometimes submiliform. They were usually enlarged at the middle, slightly tapered or rounded at the base with a short or long neck to the apex. The pileus cross section shows the uppermost portion of the cap and into the interior, structures formally known as the pileipellis and pileus trama. The pileipellis was hyaline and subgelatinous. The pileus trama mycelium was hyaline. The gill cross section shows the span across a single gill, rotated 90**°**. This cross section reveals regular to subregular lamellar trama, hymenium with pleurocystidia, pseudocystidia, basidia, basidioles and four-spored basidia. The pleurocystidia were hyaline, clavate, mucronate and sublecythiform. More basidioles than basidia can be observed in our sample, indicating slightly immature specimens. The subhymenium did not appear incrusted.

**Figure 2.**
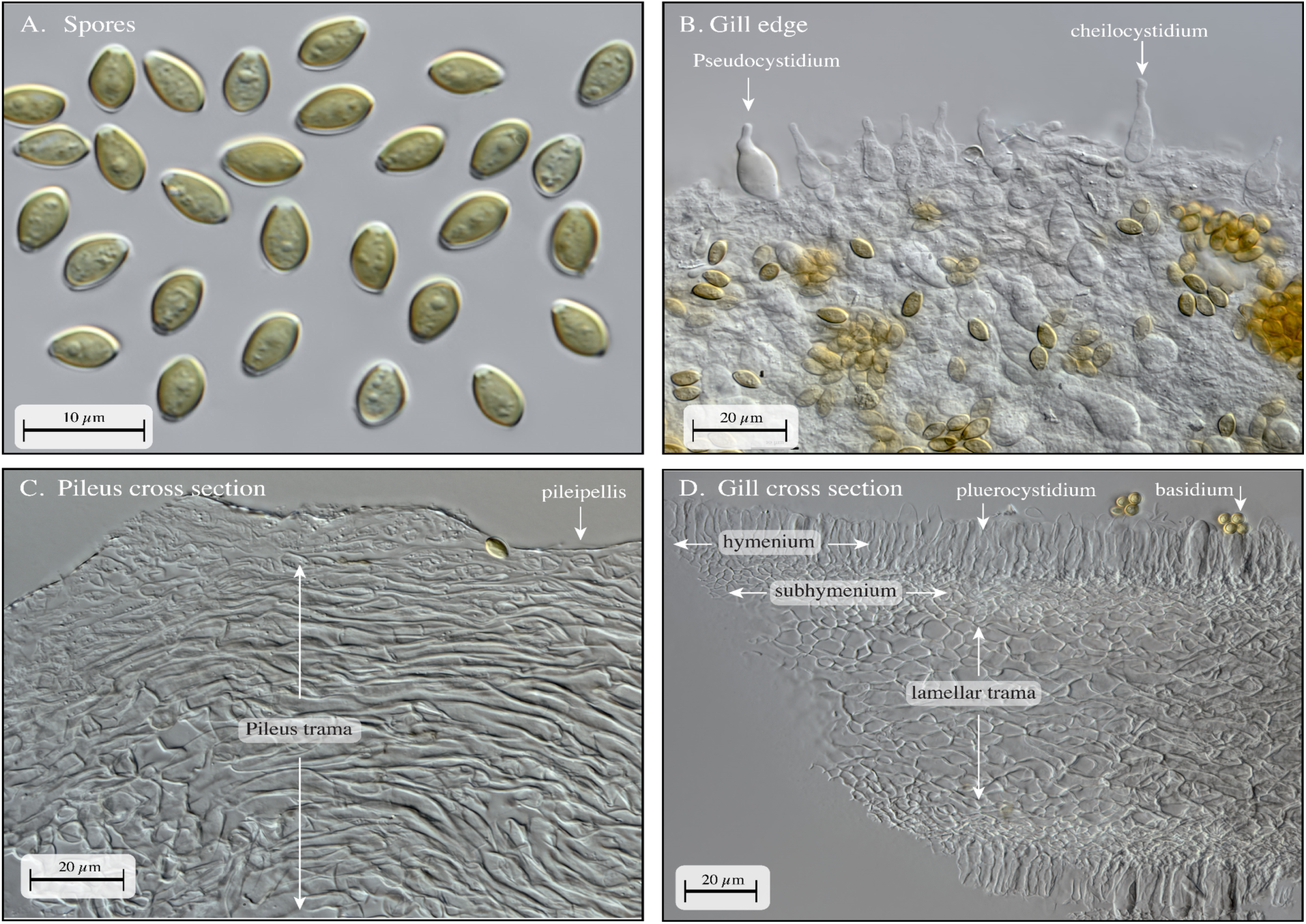
Microscopic features of *P. zapotecorum* showing. **A)** Spores, **B)** Gill edge, **C)** Pileus cross section, and **D)** Gill cross section. Dry specimens of *P. zapotecorum* were dissected, fixed, and imaged. **A)** Spores were imaged individually, and images were stitched together with Photoshop to maximize space and image clarity. Spores measured 5.9 – 6.9, averaging 6.3 ± 0.3 by 3.6 – 4.1 µm, averaging 3.8 ± 0.1, n=29. **B)** Gill edge isolated by manual manipulation and excision via forceps shows both **cheilocystidia** and the slightly more opaque **pseudocystidia** on the outer gill edge (rotated 180**°** to face upward). **C)** Pileus cross section shows the transition between the **pileus trama** and the **pileipellis**. **D)** Gill cross section shows the **hymenium** across the top and bottom both connected via the **subhymenium** to the **lamellar trama** at the core.

**Figure 3.**
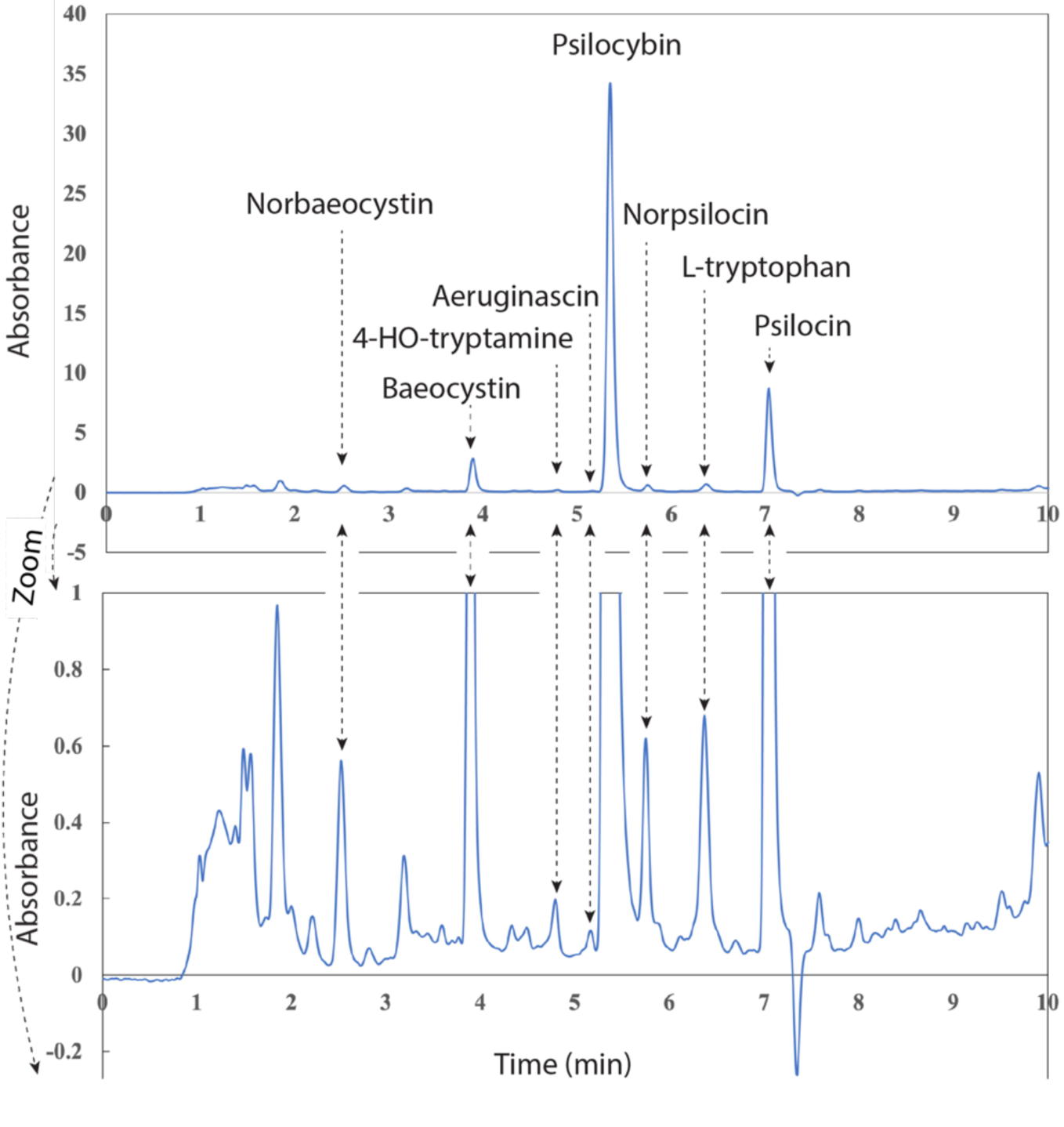
Representative chromatography profile of *P.zapotecorum* extract over *zapotecorum* extract over time. Chromatography shows absorbance at 280nm of eluant over the 10 min chromatography run, showing peaks at retention times matching that of authentic standards of norbaeocystin, standards of norbaeocystin, baeocystin, 4-HO-tryptamine, aeruginascin, psilocybin, norpsilocin, L-tryptophan, and psilocin. Separation was performed on an Thermo Accela HPLC using a phenomenex PS C18 150 x 4.6 mm 3 μm column held at 36 °C with 0.1% formic acid aqueous (mobile phase A) and 0.1% formic acid in acetonitrile (mobile phase B).

### 2.3 DNA barcoding

DNA barcoding was done to authenticate the *P. zapotecorum* used in this work. Barcoding was performed by sequencing the ITS region of the genome (**Accession: OP973765**). The Sanger sequence unambiguously assigned nucleotide base positions across the contiguous sequence, with a total read length of 667 base pairs. The complete sequence was queried using BLASTn and compared to existing records. With the results organized by percent identity and query coverage noted, we analyzed alignments.

The ITS sequence for la Martinica aligned with two other records of *P. zapotecorum.* The GenBank sequence with highest homology was a collection by Virginia Ramírez-Cruz (100% identity match, 97% query coverage, Accession: KC669303) followed by a collection from Aporo, Michoacan, Mexico in 2012, which includes field photos (99.83% identity match, 596/597 base pairs, Accession: MH050401).^16^ This latter sequence has one base reported as Y (representing either cytosine or thymine), where the sequence generated in this work contains a cytosine in that position. This cytosine lacks any ambiguity, as shown by the corresponding fluorescence peak in the electrophoretogram.

Our entry also matched with related species, albeit with lower identity. The BLAST alignment showed records of *Psilocybe subtropicalis* (accession: MH290504.1 & OM812678.1, 99.83% identity match, 89% query coverage), both with the same substitution–a thymine where a cytosine is present in *P. zapotecorum*. *Psilocybe* cf. *papuana* (accession: MH627380.1, 99.67% identity, 90% query coverage and accession: OP602367.1, 99.66% identity, 89% query coverage), and *Psilocybe keralensis* (Accession: KX357870, 99.54% identity, 98% query coverage) also matched closely. BLAST also hit a purported record of *Psilocybe caerulescens* (Genbank accession: OM276748) but published work recently showed that this record is misidentified, and is omitted from further discussion here.^63^

Section Zapotecorum was not resolvable using this one barcode. The support values within this section were all below 78. To increase the support values for resolving section Zapotecorum other genes from the full genome sequence were used.

### 2.4. Analytical chromatography

The chromatographic method resolved mixtures of psilocybin and its biosynthetic analogs with repeatable retention times (RT) [norbaeocystin (RT = 2.53 min), baeocystin (RT = 3.89 min), 4-HO-tryptamine (RT = 4.79), aeruginascin (RT = 5.16 min), psilocybin (RT = 5.35 min), norpsilocin (RT = 5.75), L-tryptophan (RT = 6.37 min), psilocin (RT = 7.04 min), and tryptamine (RT = 7.12 min), **Supplemental Figure S1]**. Partial coelutions occurred with aeruginascin and psilocybin, psilocybin and norpsilocin, and psilocin and tryptamine. The instrument response was linear with respect to analyte concentration from 0.1-100 ug/mL, allowing for a multipoint calibration curve for quantitation (SFS2). Repeat blank matrix samples spiked with low level analytes (n=10) yielded LOD and LOQ values in the single digit μg/g for five of the analytes (psilocybin, psilocin, baeocystin, norbaeocystin, and aeruginascin), in the tens of μg/g for three of the analytes (norpsilocin, tryptamine, and 4-HO-tryptamine) and in the 100s of μg/g for one analyte (L-Tryptophan) (**Table 1**). When extracts of *P. zapotecorum* fruit bodies were analyzed, the compounds of interest matching known retention times were and confirmed with UV spectra. In total, 21 samples (n= 21) were analyzed. Samples included whole mushrooms (n=7), caps (n=7), and stems (n=7).

Psilocybin was found to be the major tryptamine present. The whole fruit ranged in extracted psilocybin concentration from 10.6 to 25.7 mg/g, with a mean of 17.9 ± 1.7 mg/g (n=7). Within these sample sets psilocybin concentration was found to range from 8.9 (min concentration tested was found in the stem sample set) to 30.4 mg/g (maximum concentration tested was found in the cap sample set). On average the cap tested more potent than the stem. The stems ranged from 8.9 to 23.5 mg/g, with a mean value of 15.6 ± 2.2 mg/g (n=7). The caps ranged from 19.4 to 30.4 mg/g, with a mean value of 25.5 ± 1.8 mg/g (n=7).

Psilocin was found to be the second highest tryptamine present. Psilocin concentration in the whole fruit ranged from 0.38 to 6.52 mg/g, with a mean of 2.02 ± 0.95 mg/g (n=7). On average the stem was less potent than the cap with stems ranging from 0.30 to 2.24 mg/g, with a mean value of 1.09 ± 0.24 mg/g (n=7) and the caps ranging from 1.11 to 5.12 mg/g, with a mean value of 3.37 ± 0.53 mg/g (n=7).

Baeocystin was found to be the third highest tryptamine present, also in the mg/g range. Baeocystin concentration in the whole fruit ranged from 0.0.24 to 3.21 mg/g, with a mean of 1.34 ± 0.42 mg/g (n=7). Again, the stem was less potent than the cap with stems ranging from just under the limit of quantitation (<LOQ of 25 μg/g) to 1.17 mg/g, with a mean value of 0.57 ± 0.13 mg/g (n=7) and the caps ranging from 0.46 to 2.71 mg/g, with a mean value of 1.40 ± 0.27 mg/g (n=7).

The fourth highest tryptamine present was norbaeocystin, slipping into sub mg/g concentrations. Norbaeocystin concentration in the whole fruit ranged from 0.80 to 1.11 mg/g, with a mean of 0.50 ± 0.15 mg/g (n=7). The stem being ever less potent than the cap ranged from under the limit of quantitation (<LOQ of 22 μg/g) to 0.36 mg/g, with a mean value of 0.57 ± 0.13 mg/g (n=7) and the caps ranging from 0.46 to 2.71 mg/g, with a mean value of 1.40 ± 0.27 mg/g (n=7).

The fifth highest tryptamine present was norpsilocin. Norpsilocin concentration in the whole fruit ranged from under the limit of quantitation (<LOQ of 40 μg/g) to 1.42 mg/g, with a mean of 0.24 ± 0.20 mg/g (n=7). The stem still showed different than the cap ranging from under the limit of quantitation (<LOQ of 40 μg/g) to 70 μg/g, with a mean value of 20 ± 10 μg/g (n=7) for the stem and a range of 70 to 280 μg/g, with a mean value of 180 ± 30 μg/g for the cap(n=7).

The sixth highest tryptamine present was aeruginascin. Aeruginascin concentration in the whole fruit ranged from under the limit of quantitation (<LOQ of 21 μg/g) to 29 μg/g, with a mean of 26 ± 7 μg/g (n=7). The stem concentration ranged from under the limit of quantitation (<LOQ of 21 μg/g) to 58 μg/g, with a mean value of 32 ± 8 μg/g (n=7). The stem was less than the cap which ranged from 30 to 110 μg/g, with a mean value of 64 ± 13 μg/g (n=7).

The sixth highest tryptamine present was aeruginascin. Aeruginascin concentration in the whole fruit ranged from under the limit of quantitation (<LOQ of 21 μg/g) to 29 μg/g, with a mean of 26 ± 7 μg/g (n=7). The stem concentration ranged from under the limit of quantitation (<LOQ of 21 μg/g) to 58 μg/g, with a mean value of 32 ± 8 μg/g (n=7). The stem was less than the cap which ranged from 30 to 110 μg/g, with a mean value of 64 ± 13 μg/g (n=7).

The three analytes that were tracked with the lowest concentrations were L-tryptophan, Tryptamine and 4-OH-tryptamine. These analytes were often under the limit of quantitation. The values that were able to be quantified can be found in **Table 1**.

### 2.5 Full genome sequencing

The final *P. zapotecorum* la Martinica assembly (Accession: SAMN37305711, BioProject: PRJNA10132202) contains a total length of 59,893,331 bp across 1,849 contigs that were at least 500 bp in length. The largest contig is 2,860,434 bp with an N50 of 107,643 bp. The auN statistic, which is similar to the N50 but gives more weight to longer contigs, was 355,975, suggesting the presence of high-quality, long contigs in the assembly. The assembly had a GC content of 46.55% and no ambiguous bases (’N’s) were reported in the assembled contigs. The assembly demonstrated remarkable completeness against both Basidiomycota (n=1764; Complete: 96.6%, Single: 88.0%, Duplicate: 8.6%) and Agaricales (n=3870; Complete: 95.8%, Single: 88.5%, Duplicate: 7.3%) databases. These scores surpass many reported in literature, including the 90.3% for *Ganoderma lucidum* benchmarked against the Basidiomycota database and align closely with the highly complete genome of *Agaricus bisporus* (98.2%) benchmarked against the Agaricales database.^64,65^ The full assembly of *P. zapotecorum* La Martinica is available in the **Supplemental File 1**.

The psilocybin gene cluster was found on the 5’ end of the 71,909 bp long contig_2716 (**Figure 4**). The composition and gene order of the *P. zapotecorum* cluster is nearly identical to that of the cluster in *P. mexicana,* which is also in clade I. It contains single orthologs of PsiD, PsiK, PsiT2, and PsiM, and two paralogs of PsiH. Both clade I clusters are flanked by a hypothetical protein (hyp) gene and a 60s ribosomal protein subunit l34 gene upstream of PsiM. The hyp gene is in opposite orientation in the two species, and an additional protein of unknown function is situated between the hypothetical protein and the 60s protein genes in *P. zapotecorum*. The clade I clusters differ from the clusters in the clade II species, *P. cubensis* and *P. cyanescens,* primarily by the inclusion of PsiT1 in clade I, and the alternate positioning of PsiD. PsiD is adjacent to and divergently transcribed from PsiK in clade I species versus adjacent and transcribed in the same direction as PsiM in clade II species. The majority of the composition of the scaffolds containing the psilocybin cluster in *P. zapotecorum* and *P. Mexicana* are similar, although there is a 6kb region adjacent to PsiD of the psilocybin cluster that is not alignable and appears to be noncoding. Homologs of the laccase, PsiL, and the phosphatase, PsiP, which are involved in conversion of psilocybin to chromophoric oligomers, were also found in separate loci in *P. zapotecorum*.^24^

### 2.6 Phylogenetic analysis

The phylogenetic trees of ITS and *rpb1* reproduced expected relationships among *Psilocybe* species with notable exceptions (**Figure 5**). *Psilocybe* Clade I and Clade II were recovered as two distinct clades with high support (bp >95) in both analyses. Both trees also placed Section *Zapotecorum* within subclade A of Clade I. *P. zapotecorum* La Martinica was adjacent (bp 95,99 for ITS and *rpb1* respectively) to *P. zapotecorum* R. Heim specimen Ps-317 in section Zapotecorum.^16^ The *rpb1* gene extracted from the *P. zapotecorum* assembly was 100% identical to the reference sequence (97% query coverage). While topologies were highly congruent between the two loci in Clade I where supported (bp >80), placement of multiple species differed in Clade II with little support.

**Figure 5.**
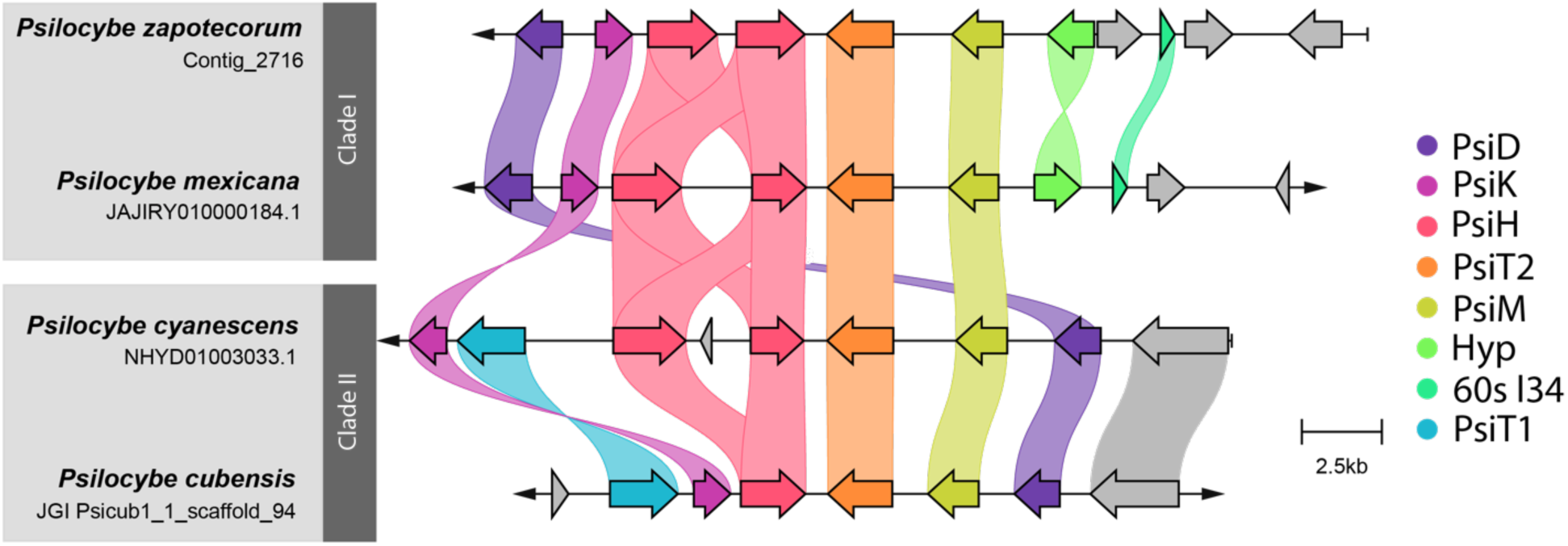
Alignment of psilocybin biosynthesis gene clusters in clade I and clade II *Psilocybe* species. Homologs of genes across species are distinguished by gene color and connected by similarly colored links. PsiD = tryptophan decarboxylase, PsiH = tryptamine-4-hydroxylase, PsiK = 4-hydroxytryptamine kinase, PsiM = N-methyltransferase, PsiT2 = psilocybin associated transporter, PsiT1 = clade II psilocybin associated transporter, 60s l34 = 60s ribosomal protein subunit l34, Hyp = hypothetical protein of unknown function. Genes in grey have limited shared synteny across species and are not known to be associated with psilocybin biosynthesis. Arrows at end of contigs indicate continuation of the contig, and bars indicate the end of the contig.

## Discussion

*P. zapotecorum* produces some of the largest and most visually striking psychedelic mushrooms found in neotropical Mexico. These mushrooms have found their way into the language, customs, and consciousness of Mexican culture in the Sierra Madre mountains of Oaxaca and beyond. A great deal of human effort has gone into understanding and expressing this mushroom mystically, artistically, and scientifically. Here, we combine field, microscopic, and molecular data for robust identification, present an effective method of cultivation, perform tryptamine analysis and provide the full genome sequence. This multidisciplinary approach adds distinction to this revered mushroom species and offers a strain for continued investigation.

Despite this mushroom’s cultural significance, a widely cultivated strain of *P. zapotecorum* had not previously been developed. Here, *P. zapotecorum* was successfully taken through a complete life cycle from spore to spore. Roger Heim’s observations in the late 1950’s indicated not all cultures would produce fruit bodies in laboratory conditions. To foster the production of fruit bodies under laboratory conditions, a wild collection was obtained (**Figure 1A**) and a multispore approach was taken. When a fruit body developed, it was isolated and cloned by removing an inner portion of the basidiome. This approach targets a strain that is both capable of fruiting and fruits quickly. If a multispore approach is used without the intermediate cloning step, the strongest growing mycelium may dominate, but may be less likely to fruit. The clonal isolate was tomentose and did not excrete pigment. Other strains on the multispore plate did excrete brown pigment into the media. It is unknown how or if this pigment affects resistance to environmental pressures, fruiting speed, or the psychedelic experience. This new fruiting strain was named ‘La Martinica’, to honor the locality from where the mushroom was collected– it is available for future investigations.

La Martinica was fruited using common materials, upcycling agricultural byproducts into medicines. In nature, *P. zapotecorum* is lignicolous, deriving nutrients from decaying woody plant matter often buried under clay. In previous studies, the substrates were formulated based on the environment where *P. zapotecorum* grows, including materials found in landslides and marshy areas such as straw, moss, compost, and sand.^4,23^ In cultivation of *P. cubensis*, it is common to use grain as spawn and a mixture of coco coir and vermiculite as substrate.^66^ Considering both standard cultivation techniques of *P. cubensis* and natural habitat of *P. zapotecorum*, grain was investigated for spawn and coco coir and vermiculite as substrate. Grain offers a nutrient dense culture and *P. zapotecorum* readily colonized oat in axenic culture. Coco coir and vermiculite as a substrate and casing layer increase air exchange and absorb and retain water. They may also already be on hand for the average mushroom grower. This composition was simple and reproducible for controlled studies. The results show that habitat-mimetic substrate is not an essential factor for cultivating *P. zapotecorum*.

Environmental parameters for cultivation were similar to that of *P. cubensis* with respect to temperature and light but deviated with regards to humidity and aeration. In the native habitats, temperatures can range from lows of ∼7°C and highs up to 29°C during mushroom season. While this wide distribution suggests that the species can tolerate varying temperatures, a modest and more tightly controlled temperature was chosen for fruiting in the laboratory. These conditions (20-22°C) mimic native average daytime high temperatures, minimize process variables, and are achievable in most laboratory settings. It was found that low intensity light was favorable and when light levels were strong, fruits would grow away from the source and sometimes abort. In the winter, fruiting in native habitats slows or stops completely. It’s not known if the lower temperatures of the winter dry season inhibit fruits or if the arid climate is the impediment. However, proper humidity control, achieved by an ultrasonic fogger, was found to be an essential factor for fruiting. These conditions are reminiscent of the low fog that blows over the cloud forests of the Sierra Madre mountains. Yet, these mountains also experience heavy rains. Perhaps the evaporation period between typical afternoon rains is important for successful fruiting. Another factor that may be important in aeration and absorbing and maintaining hydration is the top, uncolonized substrate layer. Colonized mycelium is hydrophobic, and water can pool on densely colonized surfaces. Humidifying the chamber with a spray bottle, as is typical in *P. cubensis*, caused *P. zapotecorum* fruits to abort. However, when watered by a humidification system with small, aerosolized droplets in fresh air, fruits developed prolifically. These changes in humidification, aeration and the addition of a casing layer resulted in *P. zapotecorum* fruits with morphology resembling that found in the wild.

From wild spores to a domesticated fruiting strain, the *P. zapotecorum* strain La Martinica was grown through a full life cycle yielding uniform clusters of sturdy fruits. Cultivated primordia initiated beneath uncolonized substrate, and developed into tight, caespitose clusters (**Figure 1 B**). Stems were decorated with floccose adornments, caps scalloped, finely papillate, with inrolled margins and partial veil remnants. Caps varied from umbonate, convex, or subsecotioid, but all developed and ejected spores. If left unharvested, the caps began to blacken. Sizes of spore-ejecting fruits varied but grew as tall as 33cm with caps as wide as 8cm. On the inside, the stems were dense with woody brown mycelium and a hollow center channel. When harvested, the fruits displayed immediate and intense blue bruising at points of surface trauma. The cultivated strain grew well and lives up to the reputation of being some of the largest psychoactive fungi.

Microscopic investigation of the spores and fruit bodies provide further insight and confirmation of identity. The spores were purple/brown by eye, a key feature of the genus *Psilocybe*. Under the microscope, they appeared golden and translucent (**Figure 2A**) and showed lipid droplets, indicating maturity. A unique feature in *P. zapotecorum* is the presence of pseudocystidia (**Figure 2B**).^1^ This feature helps to distinguish *P. zapotecorum* from related species such as *P. subtropicalis*. The pseudocystidia were found arising from the lamellar trama and were more opaque than other cystidia. To the untrained eye, the pseudocystidia can be difficult to distinguish from other cystidia on the surface of the hymenium. By shifting the focus while observing, one can see the origin of these cells are deeper than other cystidia on the hymenium. Crush mounting the specimen can be used to release these structures from the hymenium, where the full length and shape can be observed unobscured (not shown). Cheilocystidia were also observed and found to be polymorphous (hyaline, langeniform, ampulliform and sometimes furcate, [**Figure 2B**]). In Guzmán’s emendation, he similarly notes that the size and form of the cheilocystidia are highly variable, and he regards them as lacking taxonomic value. Cross sectional analysis of the upper most portion of the cap and into the interior (**Figure 2C**), shows the shape and organization of the hyaline mycelium in the pileus trama. In the lamellar trama, where the gill edge was broken, a single cell layer can be seen, revealing the unobstructed structure of the lamellar trama (**Figure 2D**). In the hymenium, pleurocystida, pseudocystidia, basidia and basidioles were observed. More basidioles than basidia were observed, indicating slightly immature specimens. The yellowish-brown pigment in the subhymenium described by Guzmán in 2012 was not observed.^1^ Through this microscopic analysis, the subtleties and variety discussed may aid others in identification and discernment of the pseudocystidia from pleurocystidia and cheilocystidia. Images were acquired on an Olympus BX41 microscope and images were taken using a Nikon Z7 camera. Images were further processed with Helicon Focus 8 and Photoshop.

In addition to microscopy, the DNA barcode aligns with *P. zapotecorum*. When the ITS sequence (**Accession: OP973765**) was compared to other sequences in Genbank using NCBI BLASTn, it shared the highest sequence identity with an entry by V. Ramirez-Cruz followed by one from A. Rockefeller. The entry from Ramirez-Cruz shares 100% identity, and links to a publication detailing morphology and habitat.^16^ The alignment with the Rockefeller accession may differ by a base pair - with theirs having one polymorphic read. The ambiguous read may represent a single nucleotide polymorphism within the ITS of *P. zapotecorum.*^67^ As more barcodes of *P. zapotecorum* are accessioned, this difference should become clear. Combined with habitat and morphological features, alignments with verified accessions solidify support for identification as *P. zapotecorum*.

Our entry also shared homology with species closely related to *P. zapotecorum.* Two records of *Psilocybe subtropicalis* showed high homology, both with the same substitution – a thymine for a cytosine in *P. zapotecorum.* This species shares some morphological characteristics with *P. zapotecorum,* but they can be differentiated in the field. *P. subtropicalis* has a striate subpapulate cap, a deep reddish-brown stipe with floccules appressed rather than pronounced, and a smaller stature. Its habitat also distinguishes the two. *P. subtropicalis* is found fruiting solitary or scattered in pastures, often fruiting near *Psilocybe mexicana.* It also may have a smaller distribution – *P. subtropicalis* has only been documented in the states of Veracruz and Oaxaca, whereas *P. zapotecorum* has been documented across the neotropics, from Mexico to Brazil.

This single base pair substitution suggests that *P. subtropicalis* is closely related to *P. zapotecorum*, with nearly the entire ITS region conserved between the two species yet field observations clearly resolve the two.

Next, we wanted to investigate their small molecule profile, focusing on tryptamines. A recent publication surveying an extensive collection of psychedelic mushrooms (82 collections, 31 species) reported *P. zapotecorum* to have the 5th highest psilocybin content (10 mg/g). The depth and intensity of bruising observed in *P. zapotecorum* may be a consequence of this high psilocybin content. The highest psilocybin concentration reported in the survey went to *P. serbica var. Bohemica* at 16mg/g only slightly above that of *P. azurescens* reported in other work to contain 15 ± 1 mg/g psilocybin.^67,68^ However, the *P. zapotecorum* samples in that work were analyzed two years after collection, at which point psilocybin may have degraded, decreasing concentrations.^28^ Our analysis showed higher concentration of psilocybin than any species analyzed in that survey (17.9 ± 1.7 mg/g, n=7, **Figure 4**), placing *P. zapotecorum* at the top of the potency scale. This may be a result of intrinsic psilocybin content combined with efficient sample extraction methods and HPLC analysis of samples less than six months post-harvest.^29^ The decrease in handling of laboratory grown samples compared to field collections may also preserve psilocybin content. This work places it just above P. *azurescens and P. serbica var. Bohemica* as the most potent *Psilocybe* mushroom of the world.

We wanted to understand if there was a genetic reason for the elevated psilocybin content. To answer this question, the full genome of *P. zapotecorum* was sequenced using both Illumina and Oxford Nanopore Technologies. We found that the composition and order of genes was very similar to that of *P. mexicana*, consistent with their occurrence in the same clade (**Figure 5**).^69^ Like other species with high psilocybin content, clade I species have two paralogs of PsiH, which may increase the rate of hydroxylation of tryptamine, but this has not been investigated.^18^ Different gene orientation at the 5’ end of the PsiM gene is another possible mechanism of differential pathway activity. Differential competition for promoter binding could favor the expression of *P. zapotecorum* PsiM over *P. mexicana*, which shares an upstream intergenic region with a hypothetical protein. It remains to be investigated whether any of these genes are differentially active or the enzymes they encode have differential efficiencies. It is also possible that availability of precursors or regeneration or compartmentalization lead to high concentrations of psilocybin in *P. zapotecorum*.^70–72^ Given the brilliant bluing and presence of PsiL and PsiP in this species, high psilocybin is not likely the result of reduced conversion of psilocybin to chromophoric oligomers. The sequence of the *P. zapotecorum* genome will facilitate direct investigation of all this enzymatic diversity both within and beyond the psilocybin cluster. Given the high completeness and contiguity of the *P. zapotecorum* La Martinica assembly, it is suitable for future complete gene cluster and multi-omics analysis and its development as a model system.^64,65^

Using the genome, we wanted to understand what barcodes would be sufficient for species resolution without the aid of field observations. Differentiating members within section *Zapotecorum* may not be possible with ITS analysis, due to low ITS sequence variability between its members. Using genes with higher variability, such as the *rpb1, tef-1,* or *lsu* – all protein-coding genes – a multi-locus analysis was used to build more robust branch support within section Zapotecorum. Of these genes, *rpb1* has the most information reported from other species and was therefore used in constructing a phylogenetic tree. Using both the ITS and *rpb1* sequences phylogenetic trees were constructed inferring evolutionary relationships (**Figure 6**). Both the ITS and *rpb1* trees are rooted at the junction of Clade I, which includes section Zapotecorum, and Clade II, which includes *P. cubensis* (**Figure 6AB**).^16^ Both phylogenetic trees shows section Zapotecorum as a monophyletic clade indicated by a single branch origin, which is consistent with prior work.^16^ Determining branching order within section *Zapotecorum* may not be reliable with ITS analysis, indicated by low branch support values within the clade (**Figure 6A**). However, the *rpb1*-derived tree shows high branch support within section Zapotecorum (**Figure 6B**). Notable conflict between rpb1 and ITS topologies in Clade II, with limited support, highlights the necessity of examining multiple loci to infer evolutionary relationships in *Psilocybe* as shown previously and suggests a need for further molecular systematics and/or phylogenomics in the genus.^73,74^ The utility of these and other barcode loci will become more apparent as more information is gathered on the population-level sequence variability of *P. zapotecorum* and as more species’ genomes are sequenced.

**Figure 6.**
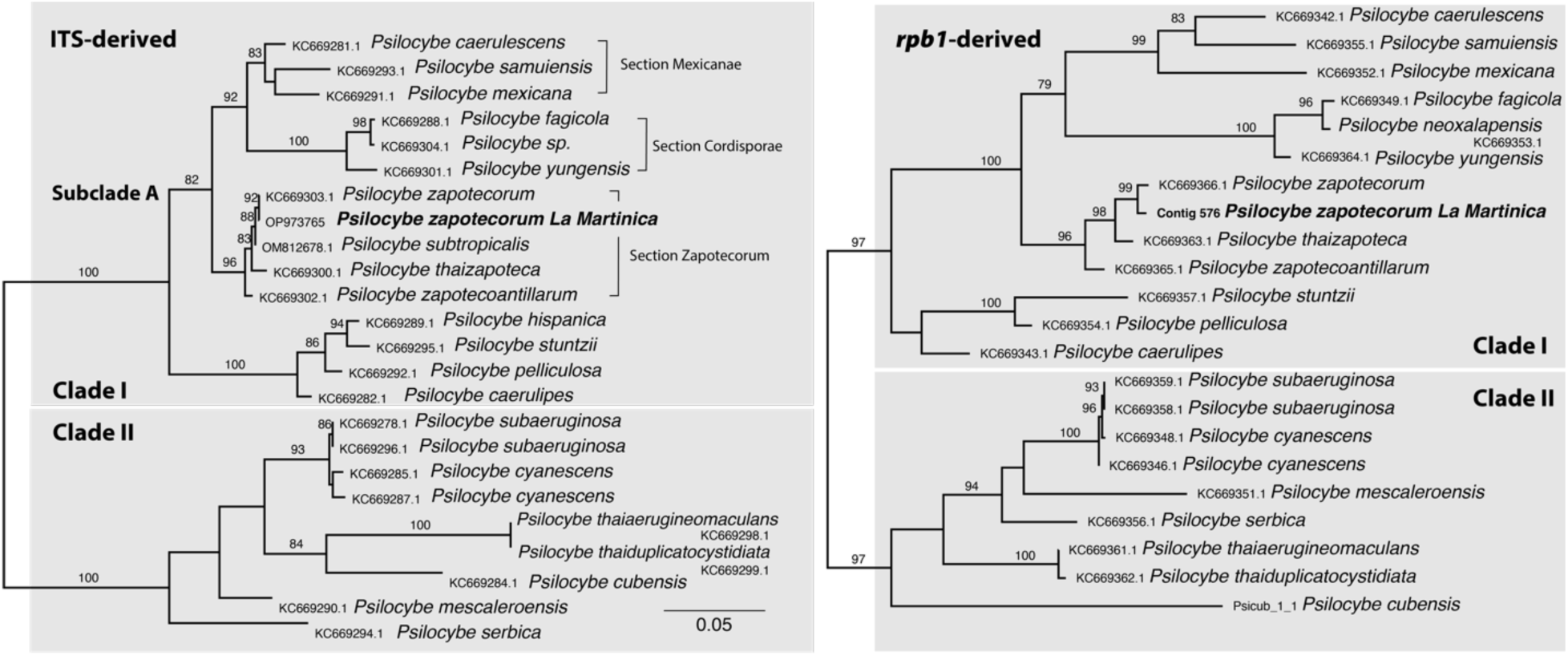
Phylogenetic placement of *P. zapotecorum* among closely related species. Phylogenetic tree was inferred from an alignment of both ITS sequences (**left panel**) and *rpb1* sequences (**right panel**). *P. zapotecorum* La Martinica (this study) is monophyletic with *P. zapotecorum* R. Heim, and the phylogeny shows *P. subtropicalis* as the adjacent taxon in the ITS tree. Clade I and Clade II, subclade A and sections Mexicanae, Cordisporae and Zapotecorum are defined as in Ramírez-Cruz et. al., 2013 and are supported in both the ITS-derived and the *rpb1*-derived tree.^16^ Support values represent the percentage of the 1000 ultrafast bootstraps in IQ-TREE. Node support values ≥ 75% bp are included. Branch lengths indicate nucleotide substitutions per site as indicated in the scale bar.

Other chemistries present may add tonal qualities to the experience of these psychedelic mushrooms. This concept is referred to as the entourage or ensemble effect and other tryptamines within psilocybin mushrooms can act synergistically with psilocybin. The other tryptamines, or minor tryptamines, include those along the biosynthetic pathway from L-tryptophan to psilocybin (4-HO-tryptamine, norbaeocystin, baeocystin, norpsilocin, and psilocin) or molecules with additional methylations (aeruginascin). These minor tryptamines are bioactive as agonists in the serotonergic pathway and may modulate the psychedelic experience. Of the six minor tryptamines resolved by our HPLC method, all six were detected [psilocin (2.02 ± 0.95 mg/g), baeocystin (1.34 ± 0.42 mg/g), norbaeocystin (0.50 ± 0.15 mg/g), norpsilocin (0.24 ±0.20 mg/g), aeruginascin (0.026 ± 0.007 mg/g), 4-HO-Tryptamine (>LOQ, 0.036 mg/g)] as well as the precursor amino acid L-tryptophan (> LOQ, 0.383 mg/g, **Figure 4**). It is still unclear how these minor tryptamines effect the psychedelic experience, if at all. The current gold standard for identification of a psychedelic is induction of a head twitch response in mice. Of the tryptamines in psilocybin mushrooms, only psilocybin and psilocin induce a head twitch response when administered individually.^75^ Co-administration of norbaeocystin and psilocybin increase head twitch response compared with psilocybin alone.^76^ However, the ratio of norbaeocystin to psilocybin in that work was not in the biological range, leaving questions as to if native concentrations affect the psychedelic experience. It is possible that the synergistic effects do not solely increase the psychedelic aspects of the mushroom experience. Some of the minor tryptamines induce other changes in mice, including thermoregulation and locomotor activity and when combined with the activity of psilocin, but may be perceived as synergistic to the psychedelic experience.^75^ Continuing to demystify the chemical composition of psychedelic plants and fungi allows for us to track experiential differences which can support safe use and personalizes medicines.

There are still many open areas of inquiry regarding the composition of these distinct mushrooms. Recently, β-carbolines were found in *P. cubensis*, *P. mexicana*, and *P. azurescens*.^46,77^ The presence of these small molecules may also act synergistically to potentiate the psychedelic experience by preventing the degradation of serotonergic drugs (monoamines) such as psilocin. It is not known if β-carbolines are present in *P. zapotecorum* and warrants further investigation. Additionally, the taste and smell of *P. zapotecorum* are unexplored. The mushroom has an electric sour taste sensation and a smell reminiscent of cucumber and radish. The sour taste has been described as similar to a 9v battery, localized to the tongue and cheeks, and lasting for at least 30 minutes, unlike typical organic acids. A cucumber odor has been tracked in other cucumber/farinaceous mushrooms to trans-2-nonenal, a component of cucumbers.^78^ Although there are no studies that we know of that investigate the aroma in *Psilocybe*, it may be part of what gives *P. zapotecorum* its cucumber radish farinaceous odor. β-carbolines may be counter-indicated with some medications and the sour sensation and distinct odor change the subjective effects immediately and profoundly. Further research into these chemistries can help understand if these components are present and if they are bioactive beyond their organoleptic properties.

Developing a basic understanding of psilocybin-containing species is a natural next step towards personalized medicine and natural product discovery. The venerable stature, intense blueing reaction, and strong sour sensation alters the subjective effects. In addition to its long history of ceremonial use by healers and shamans of southern Mexico, the symphony of pharmacological, aesthetic, and organoleptic properties seems to have set this mushroom apart for some time and this work supports the safe inclusion of *P. zapotecorum* for therapeutic use. As clinical studies of psychedelic-assisted psychotherapies gain traction, it is important to investigate the naturally occurring source of psilocybin – a wide variety of diverse mushrooms – so that synthetic psilocybin, used in most clinical studies, does not become unintentionally synonymous with mushroom therapy.

## Conclusions

As both medicine and legislation progress, questions arise as to the diversity of natural psychedelic medicines. This work touches on this complexity by focusing on a culturally relevant but rarely cultivated derrumbe mushroom, *P. zapotecorum*. The methods here detail its cultivation in common substrates. Through the process of cloning, a La Martinica strain was isolated and taken through a complete life cycle, enabling reproducible research into the chemistry and biology of this organism. Microscopy was done on the fruiting bodies showing morphological features of the cap, gills, and spores to assist in future comparisons. Chemical analysis of cultivated fruit bodies showed this strain had very high psilocybin content (17.9 ± 1.7 mg/g). Analysis of psilocybin analogs revealed modest quantities of psilocin (2.02 ± 0.95 mg/g), baeocystin (1.34 ± 0.42 mg/g), norbaeocystin (0.50 ± 0.15 mg/g), norpsilocin (0.24 ±0.20 mg/g), aeruginascin (0.026 ± 0.007 mg/g), 4-HO-Tryptamine (>LOQ, 0.036 mg/g) as well as the precursor amino acid L-tryptophan (> LOQ, 0.383 mg/g). Full genome sequencing gives insights into the psilocybin gene cluster and was used to construct a phylogenetic tree showing proposed evolutionary relationships between closely related species. This work is an interdisciplinary approach to natural product analysis in psychedelic mushrooms, incorporating mycological techniques with molecular biology and chemical testing. The demystification of these entheogenic mushrooms supports their safe use for medicinal and ceremonial purposes.

## Author contributions

D.R.M. and J.J. designed experiments, analyzed data and wrote the manuscript. H.S. designed the barcoding experiments, analyzed data and wrote the manuscript. A.R. designed the microscopy experiments and analyzed data. J.S., and I.B. helped analyze genome data and wrote the manuscript. D.E.C. oversaw experiments and wrote the manuscript.

## Acknowledgements

Special thanks to Alonzo Cortes-Perez at the Universidad de Guadalajara for the initial collection. Gratitude to Luis Talavera for sponsoring the *P. zapotecroum* genome through Entheome. Special thanks to Kelsey Scott at Ohio State University and Christopher Pauli of Tryptomics for consultation on genome assembly.

## References.

(1) Guzmán, G. New Taxonomical and Ethnomycological Observations on Psilocybe s.s. (Fungi, Basidiomycota, Agaricomycetidae, Agaricales, Strophariaceae) from Mexico, Africa and Spain. Acta botánica Mexicana 2012, 100.

(2) Heim, R. Les champignons divinatoires recueillis par Mme Valentina Pavlovna Wasson et M. R. Gordon Wasson au cours de leurs missions de 1954 et 1955 dans les pays mije, mazatèque, zapotèque et nahua du Mexique méridional et central; Gauthier-Villars]: Paris, France, 1956.

(3) Heim, R. Les Agarics Hallucinogènes Du Genre Psilocybe Recueillis Au Cours de Notre Récente Mission Dans Le Mexique Méridional et Central En Compagnie de M. R. Gordon Wasson. *Rend. Séances Acad. Sci.*, Paris 1957, 244, 695–700.

(4) Heim, R. Notes Préliminaires Sur Les Agariccs Hallucinogènes Du Mexique, IV: Breves Latinae Diagnoses Hallucinogenarum Mexicanarum Psilocybarium Ad Fera Specimina Pertinentium. Rev. Mycol 1957, 22, 58–79.

(5) Heim, R.; Cailleux, R.; Wassson, G.; Thevenard, P. Nouvelles Investigations Sur Les Champignons Hallucinogènes. Arch. Mus. Nat. Hist. Nat. 1966, 7 (9), 115–218.

(6) Heim, R.; Wasson, G. Les Champignons Hallucinogènes Du Mexique. Arch. Mus. Nat. Hist. Nat 1958, 7 (6), 1–322.

(7) Wasson, R. G. Seeking the Magic Mushroom. Life 1957, 42 (19), 100–120.

(8) Schultes, R. E. Teonanácatl: The Narcotic Mushroom of the Aztecs. American Anthropologist 1940, 42 (3), 429–443.

(9) Reko, P. B.; Groeschner, A. US National Herbarium Sheet 1745713, Barcode 00206888.

(10) Hernández-Santiago, F.; Martínez-Reyes, M.; Pérez-Moreno, J.; Mata, G. Pictographic Representation of the First Dawn and Its Association with Entheogenic Mushrooms in a 16th Century Mixtec Mesoamerican Codex. Revista mexicana de micología 2017, 46, 19– 28.

(11) Furst, P. T. Hallucinogens in Precolumbian Art; Texas Tech Press, 1974.

(12) Caso, A. Representaciones de Hongos En Los Códices. Estudios de Cultura Náhuatl 1963, No. 4, 40.

(13) Wasson, R. G. The Wondrous Mushroom: Mycolatry in Mesoamerica; McGraw-Hill, 1980.

(14) Schultes, R. E.; Bright, A. Ancient Gold Pectorals from Colombia: Mushroom Effigies? Botanical Museum Leaflets, Harvard University 1979, 27 (5/6), 113–141. www.inaturalist.org/observations/29943648.

(15) Ramírez-Cruz, V.; Guzmán, G.; Villalobos-Arámbula, A. R.; Rodríguez, A.; Matheny, P. B.; Sánchez-García, M.; Guzmán-Dávalos, L. Phylogenetic Inference and Trait Evolution of the Psychedelic Mushroom Genus Psilocybe Sensu Lato (Agaricales). Botany 2013, 91 (9), 573–591.

(16) Van Court, R. C.; Wiseman, M. S.; Meyer, K. W.; Ballhorn, D. J.; Amses, K. R.; Slot, J. C.; Dentinger, B. T. M.; Garibay-Orijel, R.; Uehling, J. K. Diversity, Biology, and History of Psilocybin-Containing Fungi: Suggestions for Research and Technological Development. Fungal Biology 2022, 126 (4), 308–319.

(17) McTaggart, A. R.; James, T. Y.; Slot, J. C.; Barlow, C.; Fechner, N.; Shuey, L. S.; Drenth, A. Genome Sequencing Progenies of Magic Mushrooms (Psilocybe Subaeruginosa) Identifies Tetrapolar Mating and Gene Duplications in the Psilocybin Pathway. Fungal Genetics and Biology 2023, 165, 103769.

(18) McTaggart, A.; McLaughlin, S.; Slot, J.; McKernan, K.; Appleyard, C.; Bartlett, T. L.; Weinert, M.; Barlow, C.; Warne, L. N.; Shuey, L. S. The Manure Tour: Invasive Populations and Clandestine Cultivars Have Bottlenecked Magic Mushrooms Since Psilocybe Cubensis Spread From Its Unknown Centre of Origin. Psilocybe cubensis.

(19) Meyer, M.; Slot, J. The Evolution and Ecology of Psilocybin in Nature. Fungal Genetics and Biology 2023, 103812.

(20) Guzman, G. The Genus Psilocybe: A Systematic Revision of the Known Species Including the History, Distribution, and Chemistry of the Hullucinogenic Species. Nova Hedwigia 1983, 74, 439.

(21) Redhead, S. A.; Moncalvo, J.-M.; Vilgalys, R.; Matheny, P. B.; Guzmán-Dávalos, L.; Guzmán, G. Proposal to Conserve the Name Psilocybe (Basidiomycota) with a Conserved Type. Taxon 2007, 56 (1), 255–257.

(22) Montiel, E.; Barragán, J. C.; Tello, I.; Mora, V. M.; León, I.; Martínez, D. Characterization and Cultivation of Psilocybe Barrerae. Micología Aplicada International 2008, 20 (2), 69–74.

(23) Lenz, C.; Wick, J.; Braga, D.; García-Altares, M.; Lackner, G.; Hertweck, C.; Gressler, M.; Hoffmeister, D. Injury-Triggered Blueing Reactions of Psilocybe “Magic” Mushrooms. Angewandte Chemie 2020, 132 (4), 1466–1470.

(24) Lenz, C.; Dörner, S.; Sherwood, A.; Hoffmeister, D. Structure Elucidation and Spectroscopic Analysis of Chromophores Produced by Oxidative Psilocin Dimerization. Chemistry–A European Journal 2021, 27 (47), 12166–12171.

(25) Stijve, T.; Meijer, A. A. R. Macromycetes from the State of Parana, Brazil. 4. The Psychoactive Species. Arquivos de biologia e tecnologia 1993, 36 (2), 313–329.

(26) Heim, R.; Hofmann, A. La Psilocybine et La Psilocine Chez Les Psilocybe et Strophaires Hallucinogenes. Les Champignons Hallucinogenes du Mexique 1958, 258–262.

(27) Gotvaldová, K.; Hájková, K.; Borovička, J.; Jurok, R.; Cihlářová, P.; Kuchař, M. Stability of Psilocybin and Its Four Analogs in the Biomass of the Psychotropic Mushroom Psilocybe Cubensis. Drug testing and analysis 2021, 13 (2), 439–446.

(28) Lenz, C.; Wick, J.; Hoffmeister, D. Identification of Ømega-N-Methyl-4-Hydroxytryptamine (Norpsilocin) as a Psilocybe Natural Product. Journal of natural products 2017, 80 (10), 2835–2838.

(29) Bigwood, J.; Beug, M. W. Variation of Psilocybin and Psilocin Levels with Repeated Flushes (Harvests) of Mature Sporocarps of Psilocybe Cubensis (Earle) Singer. Journal of Ethnopharmacology 1982, 5 (3), 287–291. 10.1016/0378-8741(82)90014-9.

(30) Carhart-Harris, R. L.; Erritzoe, D.; Williams, T.; Stone, J. M.; Reed, L. J.; Colasanti, A.; Tyacke, R. J.; Leech, R.; Malizia, A. L.; Murphy, K.; Hobden, P.; Evans, J.; Feilding, A.; Wise, R. G.; Nutt, D. J. Neural Correlates of the Psychedelic State as Determined by FMRI Studies with Psilocybin. Proceedings of the National Academy of Sciences 2012, 109 (6), 2138–2143. 10.1073/pnas.1119598109.

(31) Nayak, S. M.; Singh, M.; Yaden, D. B.; Griffiths, R. R. Belief Changes Associated with Psychedelic Use. Journal of Psychopharmacology 2023, 37 (1), 80–92.

(32) Griffiths, R. R.; Richards, W. A.; McCann, U.; Jesse, R. Psilocybin Can Occasion Mystical-Type Experiences Having Substantial and Sustained Personal Meaning and Spiritual Significance. Psychopharmacology 2006, 187, 268–283.

(33) Barrett, F.; Griffiths, R. Validation of the Revised Mystical Experience Questionnaire in Experimental Sessions with Psilocybin. J Psychopharmacol. 2015, 11, 1182–1190. 10.1177/0269881115609019.

(34) Goodwin, G. M.; Aaronson, S. T.; Alvarez, O.; Atli, M.; Bennett, J. C.; Croal, M.; DeBattista, C.; Dunlop, B. W.; Feifel, D.; Hellerstein, D. J. Single-Dose Psilocybin for a Treatment-Resistant Episode of Major Depression: Impact on Patient-Reported Depression Severity, Anxiety, Function, and Quality of Life. Journal of Affective Disorders 2023, 327, 120–127.

(35) Griffiths, R. R.; Johnson, M. W.; Carducci, M. A.; Umbricht, A.; Richards, W. A.; Richards, B. D.; Cosimano, M. P.; Klinedinst, M. A. Psilocybin Produces Substantial and Sustained Decreases in Depression and Anxiety in Patients with Life-Threatening Cancer: A Randomized Double-Blind Trial. Journal of psychopharmacology 2016, 30 (12), 1181–1197.

(36) Grob, C. S.; Danforth, A. L.; Chopra, G. S.; Hagerty, M.; McKay, C. R.; Halberstadt, A. L.; Greer, G. R. Pilot Study of Psilocybin Treatment for Anxiety in Patients with Advanced-Stage Cancer. Archives of general psychiatry 2011, 68 (1), 71–78.

(37) Bogenschutz, M. P.; Ross, S.; Bhatt, S.; Baron, T.; Forcehimes, A. A.; Laska, E.; Mennenga, S. E.; O’Donnell, K.; Owens, L. T.; Podrebarac, S. Percentage of Heavy Drinking Days Following Psilocybin-Assisted Psychotherapy vs Placebo in the Treatment of Adult Patients with Alcohol Use Disorder: A Randomized Clinical Trial. JAMA psychiatry 2022, 79 (10), 953–962.

(38) Feduccia, A.; Agin-Liebes, G.; Price, C. M.; Grinsell, N.; Paradise, S.; Rabin, D. M. The Need for Establishing Best Practices and Gold Standards in Psychedelic Medicine. Journal of Affective Disorders 2023, 332, 47–54.

(39) Greenway, K. T.; Garel, N.; Jerome, L.; Feduccia, A. A. Integrating Psychotherapy and Psychopharmacology: Psychedelic-Assisted Psychotherapy and Other Combined Treatments. Expert Review of Clinical Pharmacology 2020, 13 (6), 655–670.

(40) Kavalali, E. T.; Monteggia, L. M. Rapid Homeostatic Plasticity and Neuropsychiatric Therapeutics. Neuropsychopharmacology 2023, 48 (1), 54–60.

(41) Kim, J.-W.; Suzuki, K.; Kavalali, E. T.; Monteggia, L. M. Bridging Rapid and Sustained Antidepressant Effects of Ketamine. Trends in Molecular Medicine 2023.

(42) Moliner, R.; Girych, M.; Brunello, C. A.; Kovaleva, V.; Biojone, C.; Enkavi, G.; Antenucci, L.; Kot, E. F.; Goncharuk, S. A.; Kaurinkoski, K. Psychedelics Promote Plasticity by Directly Binding to BDNF Receptor TrkB. Nature Neuroscience 2023, 26 (6), 1032–1041.

(43) Stamets, P. Psilocybin Mushrooms of the World: An Identification Guide; Ten Speed Press, 2023.

(44) Abbas, A. I.; Carter, A.; Jeanne, T.; Knox, R.; Korthuis, P. T.; Hamade, A.; Stauffer, C.; Uehling, J. Oregon Psilocybin Advisory Board Rapid Evidence Review and Recommendations; Oregon Psilocybin Advisory Board Salem, OR, USA, 2021.

(45) Dörner, S.; Rogge, K.; Fricke, J.; Schäfer, T.; Wurlitzer, J. M.; Gressler, M.; Pham, D. N. K.; Manke, D. R.; Chadeayne, A. R.; Hoffmeister, D. Genetic Survey of Psilocybe Natural Products. ChemBioChem 2022, 23 (14), e202200249. 10.1002/cbic.202200249.

(46) Kolmogorov, M.; Yuan, J.; Lin, Y.; Pevzner, P. A. Assembly of Long, Error-Prone Reads Using Repeat Graphs. Nature biotechnology 2019, 37 (5), 540–546.

(47) Li, H.; Durbin, R. Fast and Accurate Long-Read Alignment with Burrows–Wheeler Transform. Bioinformatics 2010, 26 (5), 589–595.

(48) Bolger, A. M.; Lohse, M.; Usadel, B. Trimmomatic: A Flexible Trimmer for Illumina Sequence Data. Bioinformatics 2014, 30 (15), 2114–2120.

(49) Andrews, K. R.; Seaborn, T.; Egan, J. P.; Fagnan, M. W.; New, D. D.; Chen, Z.; Hohenlohe, P. A.; Waits, L. P.; Caudill, C. C.; Narum, S. R. Whole Genome Resequencing Identifies Local Adaptation Associated with Environmental Variation for Redband Trout. Molecular Ecology 2023, 32 (4), 800–818.

(50) Li, H.; Handsaker, B.; Wysoker, A.; Fennell, T.; Ruan, J.; Homer, N.; Marth, G.; Abecasis, G.; Durbin, R.; Subgroup, 1000 Genome Project Data Processing. The Sequence Alignment/Map Format and SAMtools. bioinformatics 2009, 25 (16), 2078–2079.

(51) Walker, B. J.; Abeel, T.; Shea, T.; Priest, M.; Abouelliel, A.; Sakthikumar, S.; Cuomo, C. A.; Zeng, Q.; Wortman, J.; Young, S. K. Pilon: An Integrated Tool for Comprehensive Microbial Variant Detection and Genome Assembly Improvement. PloS one 2014, 9 (11), e112963.

(52) Gurevich, A.; Saveliev, V.; Vyahhi, N.; Tesler, G. QUAST: Quality Assessment Tool for Genome Assemblies. Bioinformatics 2013, 29 (8), 1072–1075.

(53) Seppey, M.; Manni, M.; Zdobnov, E. M. BUSCO: Assessing Genome Assembly and Annotation Completeness. Gene prediction: methods and protocols 2019, 227–245.

(54) Altschul, S. F.; Gish, W.; Miller, W.; Myers, E. W.; Lipman, D. J. Basic Local Alignment Search Tool. Journal of molecular biology 1990, 215 (3), 403–410.

(55) Slater, G. S. C.; Birney, E. Automated Generation of Heuristics for Biological Sequence Comparison. BMC bioinformatics 2005, 6, 1–11.

(56) Gilchrist, C. L. M.; Chooi, Y. H. Automatic Generation of Gene Cluster Comparison Figures. Bioinformatics. *doi* 2020, 10.

(57) Edgar, R. C. MUSCLE: Multiple Sequence Alignment with High Accuracy and High Throughput. Nucleic Acids Res 2004, 32 (5), 1792–1797.

(58) Edgar, R. C. MUSCLE: A Multiple Sequence Alignment Method with Reduced Time and Space Complexity. BMC bioinformatics 2004, 5 (1), 1–19.

(59) Madeira, F.; Pearce, M.; Tivey, A. R.; Basutkar, P.; Lee, J.; Edbali, O.; Madhusoodanan, N.; Kolesnikov, A.; Lopez, R. Search and Sequence Analysis Tools Services from EMBL-EBI in 2022. Nucleic acids research 2022, 50 (W1), W276–W279.

(60) Nguyen, L.-T.; Schmidt, H. A.; Von Haeseler, A.; Minh, B. Q. IQ-TREE: A Fast and Effective Stochastic Algorithm for Estimating Maximum-Likelihood Phylogenies. Molecular biology and evolution 2015, 32 (1), 268–274.

(61) Hoang, D. T.; Chernomor, O.; Von Haeseler, A.; Minh, B. Q.; Vinh, L. S. UFBoot2: Improving the Ultrafast Bootstrap Approximation. Molecular biology and evolution 2018, 35 (2), 518–522.

(62) Bradshaw, A. J.; Backman, T. A.; Ramírez-Cruz, V.; Forrister, D. L.; Winter, J. M.; Guzmán-Dávalos, L.; Furci, G.; Stamets, P.; Dentinger, B. T. DNA Authentication and Chemical Analysis of Psilocybe Mushrooms Reveal Widespread Misdeterminations in Fungaria and Inconsistencies in Metabolites. Applied and Environmental Microbiology 2022, 88 (24), e01498–22. 10.1128/aem.01498-22.

(63) Morin, E.; Kohler, A.; Baker, A. R.; Foulongne-Oriol, M.; Lombard, V.; Nagye, L. G.; Ohm, R. A.; Patyshakuliyeva, A.; Brun, A.; Aerts, A. L. Genome Sequence of the Button Mushroom Agaricus Bisporus Reveals Mechanisms Governing Adaptation to a Humic-Rich Ecological Niche. Proceedings of the National Academy of Sciences 2012, 109 (43), 17501–17506.

(64) Chen, S.; Xu, J.; Liu, C.; Zhu, Y.; Nelson, D. R.; Zhou, S.; Li, C.; Wang, L.; Guo, X.; Sun, Y. Genome Sequence of the Model Medicinal Mushroom Ganoderma Lucidum. Nature communications 2012, 3 (1), 913.

(65) McCoy, P. Radical Mycology: A Treatise on Seeing and Working with Fungi; Chthaeus press Portland, OR, USA, 2016.

(66) Gotvaldová, K.; Borovička, J.; Hájková, K.; Cihlářová, P.; Rockefeller, A.; Kuchař, M. Extensive Collection of Psychotropic Mushrooms with Determination of Their Tryptamine Alkaloids. International Journal of Molecular Sciences 2022, 23 (22), 14068. 10.3390/ijms232214068.

(67) Stamets, P.; Gartz, J. A New Caerulescent Psilocybe from the Pacific Coast of Northwestern America. Integration 1995, 6, 21–28.

(68) Rokas, A.; Mead, M. E.; Steenwyk, J. L.; Raja, H. A.; Oberlies, N. H. Biosynthetic Gene Clusters and the Evolution of Fungal Chemodiversity. Natural product reports 2020, 37 (7), 868–878.

(69) Lenz, C.; Sherwood, A.; Kargbo, R.; Hoffmeister, D. Taking Different Roads: L-Tryptophan as the Origin of Psilocybe Natural Products. ChemPlusChem 2021, 86 (1), 28–35.

(70) Fricke, J.; Blei, F.; Hoffmeister, D. Enzymatic Synthesis of Psilocybin. Angewandte Chemie International Edition 2017, 56 (40), 12352–12355.

(71) Blei, F.; Fricke, J.; Wick, J.; Slot, J. C.; Hoffmeister, D. Iterative L-Tryptophan Methylation in Psilocybe Evolved by Subdomain Duplication. ChemBioChem 2018, 19 (20), 2160–2166.

(72) Bradshaw, A. J.; Ramirez-Cruz, V.; Awan, A. R.; Furci, G.; Guzman-Davalos, L.; Stamets, P.; Dentinger, B. T. Phylogenomics of the Psychoactive Mushroom Genus Psilocybe and Evolution of the Psilocybin Biosynthetic Gene Cluster. bioRxiv 2022, 2022–12.

(73) Borovička, J.; Oborník, M.; Stříbrnỳ, J.; Noordeloos, M. E.; Sánchez, L. A.; Gryndler, M. Phylogenetic and Chemical Studies in the Potential Psychotropic Species Complex of Psilocybe Atrobrunnea with Taxonomic and Nomenclatural Notes. Persoonia-Molecular Phylogeny and Evolution of Fungi 2015, 34 (1), 1–9.

(74) Glatfelter, G. C.; Pottie, E.; Partilla, J. S.; Sherwood, A. M.; Kaylo, K.; Pham, D. N. K.; Naeem, M.; Sammeta, V. R.; DeBoer, S.; Golen, J. A.; Hulley, E. B.; Stove, C. P.; Chadeayne, A. R.; Manke, D. R.; Baumann, M. H. Structure–Activity Relationships for Psilocybin, Baeocystin, Aeruginascin, and Related Analogues to Produce Pharmacological Effects in Mice. ACS Pharmacol. Transl. Sci. 2022, 5 (11), 1181–1196. 10.1021/acsptsci.2c00177.

(75) Anas, N. A. The Effects of E. Coli Derived Psilocybin on the Gut Microbiome. PhD Thesis, Miami University, 2022.

(76) Blei, F.; Dörner, S.; Fricke, J.; Baldeweg, F.; Trottmann, F.; Komor, A.; Meyer, F.; Hertweck, C.; Hoffmeister, D. Simultaneous Production of Psilocybin and a Cocktail of β-Carboline Monoamine Oxidase Inhibitors in “Magic” Mushrooms. Chemistry–A European Journal 2020, 26 (3), 729–734.

(77) Wood, W. F.; Brandes, M. L.; Watson, R. L.; Jones, R. L.; Largent, D. L. Trans-2-Nonenal, the Cucumber Odor of Mushrooms. Mycologia 1994, 86 (4), 561–563.

