## Supplemental files for "Cultivation, chemistry, and genome of *Psilocybe zapotecorum*"

### Supplemental Figures

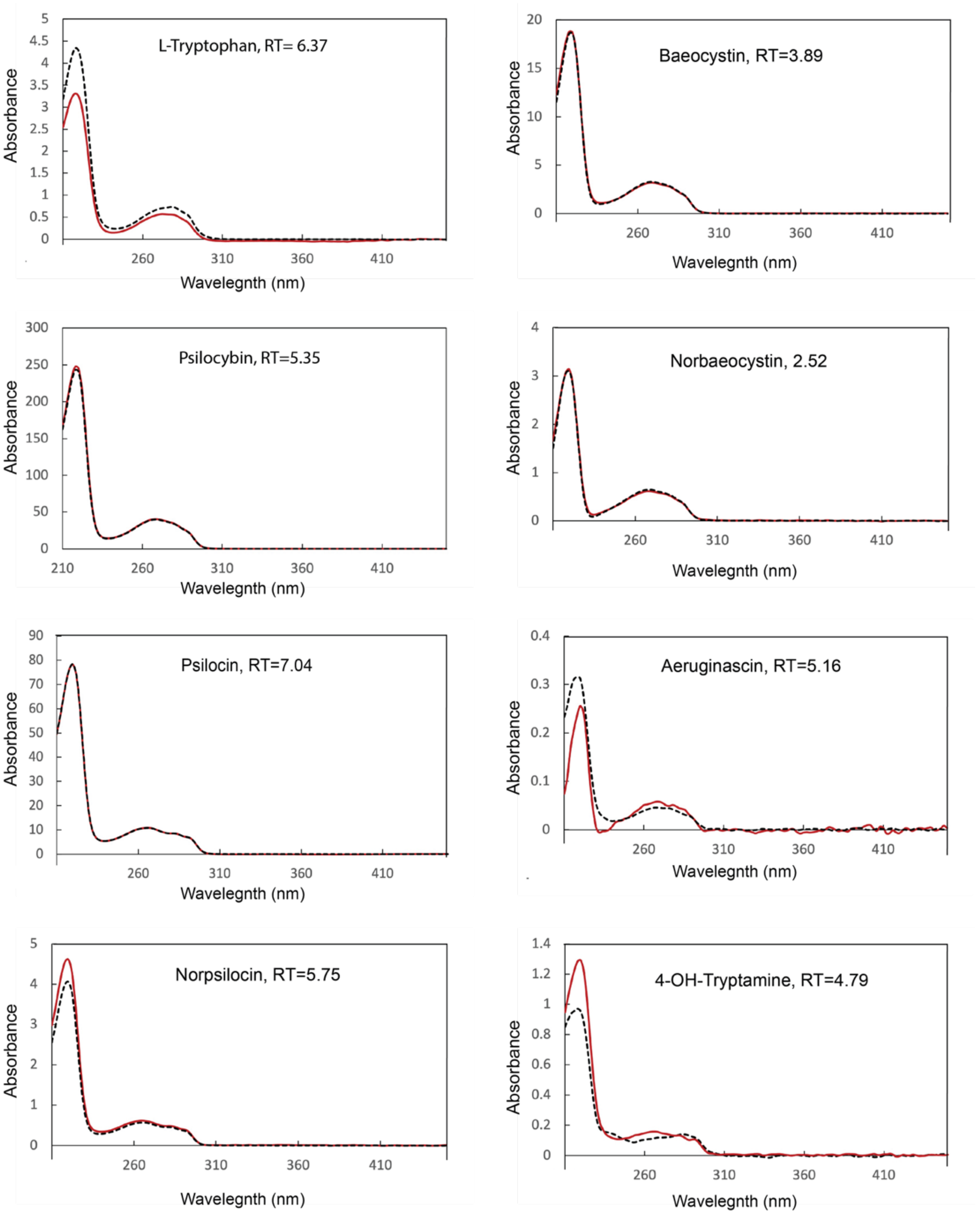


**Supplemental Figure 1.** Representative UV spectra from 200-460nm of CRM standards (black dotted curve) overlaid with peaks from *P. zapotecorum* extract (red curve) at the noted retention times. Standards were run at a concentration similar to that found in the mushroom extract. The standard runs were also transformed to match the concentration of the samples so that the shapes of the curves could be more easily compared.
